# Interactions between mechanisms of reproductive isolation

**DOI:** 10.1101/2025.02.20.639255

**Authors:** Alexandre Blanckaert, Vitor C. Sousa

## Abstract

Speciation is responsible for the diversity of species observed today and corresponds to the build-up of reproductive isolation between populations. Reproductive isolation can be generated by different mechanisms that have been extensively characterized, yet how their interactions affect speciation remains largely unknown. Here, we explicitly model the interaction of three key mechanisms (local adaptation, mate choice and genetic hybrid incompatibilities) quantifying their relative contribution to the evolution of reproductive isolation. We modeled two populations exchanging migrants using Fisher’s Geometric Model for local adaptation, phenotype matching for mate choice, and multiple pairs of Bateson-Dobzhansky-Muller Incompatibilities (DMI). All three mechanisms were determined by the same set of loci, creating conditions for interactions between barriers both at the genetic and population levels. We found very few cases where the three barriers evolved together. Instead, two barriers could evolve depending on the migration rate: either local adaptation and genetic incompatibilities for limited migration, or local adaptation and mate choice for higher migration. Our results showed that local adaptation due to ecological differentiation was the first to evolve and by far the most effective reproductive barrier. Finally, we demonstrated that in a polygenic model, populations could become locally adapted and evolve strict mate choice, yet they would not accumulate incompatibilities provided that there was sufficient gene flow.

---

Speciation has been a central theme in evolutionary biology since its early origin (Darwin, 1859). Yet, despite a truly large body of work (Kirkpatrick and Ravigné, 2002), general rules of how speciation works and which mechanisms are involved remain to a large extent a mystery. So far, most findings provide a case by case set of rules rather than universal ones (see Table 1 in Sobel et al., 2010). Two key questions have been under active research and debate. First, what should be measured to capture progress in the speciation process (Westram et al., 2022; Mallet and Mullen, 2022)? Second, what is the end of the speciation process? This was historically defined as complete reproductive isolation (RI) between diverging populations (Coyne and Orr, 2004). However, recent advances in genomics reveal that gene flow between well-defined species is far from uncommon (Payseur and Rieseberg, 2016), making it difficult to draw the line of when speciation completes (Hey and Pinho, 2012; Dopman et al., 2024). Furthermore, it has been recently argued that partial RI may not be a transient state on the way to speciation but that it may also be “a final equilibrium” state (Servedio and Hermisson, 2020; Barraclough, 2024) further blurring the notion of species.

**Table 1:** List of parameters and their default values. Status indicates whether the parameter was always constant across all simulations (Fixed), could change between simulations (Variable) or within simulations (Evolvable).

| Parameter | Variable | Default value | Status |
| --- | --- | --- | --- |
| Population size | $N$ | 10,000 | Variable |
| Number of loci | $L$ | 500 | Variable |
| Number of linkage blocks | $n_{LB}$ | 5 | Fixed |
| Recombination rate between adjacent loci<br>(per individual per generation) | $\rho$ | 0.001 | Fixed |
| Recombination rate between linkage block<br>(per individual per generation) | $\rho_{LB}$ | 0.01 | Fixed |
| Recombination rate between choosiness locus and first locus<br>(per individual per generation) | $\rho_{MC}$ | 0.1 | Fixed |
| Number of phenotypic traits | $n_t$ | 1 | Variable |
| Mutation rate (per individual per base pair per generation) | $\mu$ | $10^{-6}$ | Fixed |
| Standard deviation of mutational effects | $\sigma_\mu$ | 0.05 | Fixed |
| Standard deviation for mutation at the choosiness loci | $\sigma_P$ | 0.05 | Fixed |
| Standard deviation for mutation at the cost loci | $\sigma_C$ | 0 | Fixed |
| Shape of Fisher's Geometric model<br>controlling the average epistasis | $Q$ | 2 | Variable |
| Optimal phenotype at trait $j$ for environment A (and B) | $z_j^A$ ($z_j^B$) | 5 (-5) | Variable |
| Squared distance between perfectly adapted individuals<br>from environment A and B | $\Delta_{AB}$ | 100 | Variable |
| Fitness of an optimal phenotype for environment B (A)<br>in the A (B) environment | $\Omega^B$ ($\Omega^A$) | 0.1 | Variable |
| Probability for a male perfectly adapted to environment A (resp. B)<br>to accepting a female perfectly adapted to environment B (resp. A) | $P_C$ | 1 | Evolvable |
| Number of potential partner to choose among | $\chi$ | 100 | Variable |
| Number of pairwise DMIs | $n_{DMI}$ | 50 | Variable |
| Epistasis due to DMIs | $\epsilon$ | 0.1 | Variable |
| Migration rate | $m$ | 0.05 | Variable |

Different mechanisms of RI can drive the speciation process. They have been classified depending on the timing (pre- or postzygotic) and nature of selection acting upon the mechanism of RI (extrinsic or intrinsic) (Coyne and Orr, 2004). Disentangling which RI mechanism(es) have the largest impact during speciation, and whether their interactions are synergistic or antagonistic remained a central question of speciation research. In other words, do interactions lead to coupling (“coincidences of barrier effects, resulting in a stronger overall barrier to gene flow”, sensu Butlin and Smadja, 2018, synergistic interactions) or to a reduction of the overall strength of the RI barrier (antagonistic)?

There have been many attempts to characterize the joint evolution of different mechanisms of RI (e.g., mate choice, hybrid load), or the evolution of one conditioned on the other, with assortative mating models via “magic trait” or reinforcement being known examples. Indeed, the coupling of RI barriers is considered an important step in the speciation process (Barton, 1983; Butlin and Smadja, 2018; Dopman et al., 2024). Yet, it may lead to overall weaker rather than stronger RI (Aubier et al., 2024), and it remains unclear under which conditions multiple barriers lead to synergistic or antagonistic outcomes. Theoretical studies have shown that multiple RI barriers do not always act synergistically (e.g., Maisonneuve et al., 2024).

There has been a recent push to fully characterize the different RI barriers acting between pairs of nascent species. For example, male mate choice and (almost complete) hybrid sterility have been reported between *Tetranychus cinnabarinnus* and *T. urticae* spider mites (Cruz et al., 2024). Similarly, natural populations of *Xiphophorus birchmanni* and *X. malinche* swordtail fish adapted to different thermal environments have strong assortative mating (female choice). Their hybrids display “widespread misregulation” of thermotolerance genes, suggesting an overlap between genes under local adaptation and the ones responsible for genetic incompatibilities (Payne et al., 2024). However, quantifying the role of these mechanisms can be challenging, even for species amenable to laboratory manipulation (e.g., Aguillon et al., 2025). In addition, there are no general rules emerging. For example, Castillo et al. (2015) conducted experimental evolution of *Caenorhabditis remanei* on different substrates, measuring the evolution of local adaptation, mate choice and genetic incompatibilities, concluding that local adaptation was not relevant for the evolution of RI. This contrasts with other studies highlighting the importance of local adaptation in natural populations (reviewed in Table 1 of Rundle and Nosil, 2005), with many empirical examples of ecological speciation: Darwin finches (Grant and Grant, 2024), ciclids (McGee et al., 2020), freshwater stickleback (Hatfield and Schluter, 1999; Marques et al., 2019), whitefish (Rogers and Bernatchez, 2007) *Heliconius* butterfly (Mallet and Barton, 1989), and pea aphids (Via et al., 2000).

While the spatial context of speciation (i.e., allopatric, parapatric and sympatric) has historically been strongly debated, several authors proposed to consider whether speciation occurs with or without gene flow, irrespective of the spatial context (Butlin et al., 2008; Stankowski and Ravinet, 2021; Bolnick et al., 2023). When there is gene flow between populations (non sterile F_1_s), genetic incompatibilities create a hybrid load that may act as a sieve, leading to removal of genetic incompatibilities from both populations. Thus, provided that the mutations involved in incompatibilities do not confer a local selective advantage, or are not tightly linked to a mutation that does, we expect genetic incompatibilities to be lost when there is gene flow (Agrawal et al., 2011; Bank et al., 2012; Blanckaert and Hermisson, 2018). In addition, migration between two populations in contact with each other creates divergent selection on mate choice traits (McPeek and Gavrilets, 2006; Yamaguchi and Iwasa, 2013) to avoid the production of unfit hybrids (whether the cause is intrinsic or extrinsic). Overall, this makes migration and the strength of gene flow a key parameter to any study relative to speciation.

Here, we characterized the joint evolution of three specific mechanisms of RI, determining whether there were synergistic or antagonistic interactions between the different RI barriers. We investigated whether the evolution of one mechanism impacted the evolution of the others, and quantified their interaction by assessing their joint effect on RI, compared to their individual contributions. The three mechanisms of RI were chosen to represent each a different type: mate choice (here an intrinsic prezygotic mechanism), accumulation of genetic incompatibilities (intrinsic postzygotic) and local adaptation (extrinsic pre- and postzygotic). We decided to investigate this question, using a model where all three RI mechanisms interact with each other at the genetic level. By considering a common underlying genetic architecture, we expected this setup to favor the joint evolution of RI barriers, increasing the likelihood of observing interactions between barriers.

In details, we assumed a polygenic basis for adaptation (e.g., adaptation to host plants of *T. urticae* (Villacis-Perez et al., 2024); *see also (Mackay, 2001)), with the phenotype of an individual defined by the contribution of many loci. A subset of these loci formed intrinsic genetic incompatibilities, as such shared genetic basis should be common (Schluter and Conte, 2009; Kulmuni and Westram, 2017) and such pattern has been identified in Mimulus guttatus* (copper tolerance, Macnair and Christie, 1983) or *Arabidopsis thaliana* (resistance against biotrophic pathogens, Świadek et al., 2017). Recently, Frayer et al. (2025) compiled a list of known genetic incompatibilities - among them the most likely explanation for 18 of DMIs was host pathogen conflicts and adaptation for 4.2%. Moreover, the fitness of an individual is defined as the compound effect of extrinsic and intrinsic fitness; the first term is determined by the phenotype (i.e., how well adapted it is to the environment) and the second by the intrinsic genetic load due to genetic incompatibilities. Finally, (intrinsic) mate choice is based on the phenotype distance between potential mates (phenotype matching). This last assumption corresponds to a “classical magic trait” (according to Servedio et al., 2011) and has been reported in various organisms: wing pattern in *Heliconius* butterfly (Mallet and Barton, 1989; Jiggins et al., 2001, 2005) or body size in stickleback fishes (Snyder and Dingle, 1989; McKinnon et al., 2004); for a more complete list of biological putative examples, see Table 1 of Servedio et al. (2011). As argued by Thibert-Plante and Gavrilets (2013), these traits might be more common than expected, as selection favors their formation when initially absent. Overall, our approach included all five major, important features of speciation models, as identified by Kirkpatrick and Ravigné (2002). Under this model, we showed that local adaptation was the first mechanism to evolve and also the key mechanism responsible for the strength of reproductive isolation. Typically, depending on the level of migration between populations, either mate choice or the accumulation of genetic incompatibilities evolved in tandem with local adaptation. Interestingly, we found that even populations that evolve local adaptation and strict mate choice, might not accumulate genetic incompatibilities, provided that there is sufficient gene flow.

## Model

To understand the evolution of different types of RI barriers and their interactions, we considered a model with a common architecture. We investigated a polygenic model where loci determining the phenotype, which affected local adaptation and mate choice, could also be involved in genetic incompatibilities, as illustrated in Figure 1. Due to the complex polygenic genetic basis of the phenotype and genetic incompatibilities, we used individual-based simulations. To investigate the long term evolution, we followed > 10*N* generations of two populations of *N* diploid dioecious individuals exchanging migrants at rate *m* (two-island-island model). To model local adaptation, we considered that the two islands represent two environments (A and B) with different phenotypic optima, and for simplicity, we referred to the population residing in each environment as population A and B, respectively. We let the phenotype, genetic incompatibilities and mate choice evolve, exploring the impact of key processes in the build-up of RI. As detailed below, we varied: (1) the extent of migration, (2) the genetic architecture of phenotypes and intrinsic genetic incompatibilities, (3) life history of species (juvenile vs adult dispersal stage, as they can affect evolutionary outcomes (Johst and Brandl, 1997; Débarre and Gandon, 2011)), and (4) initial levels of divergence between populations (ancestral vs secondary contact), keeping the other parameters fixed (see Table 1 for the complete list of parameters).

**Figure 1:**
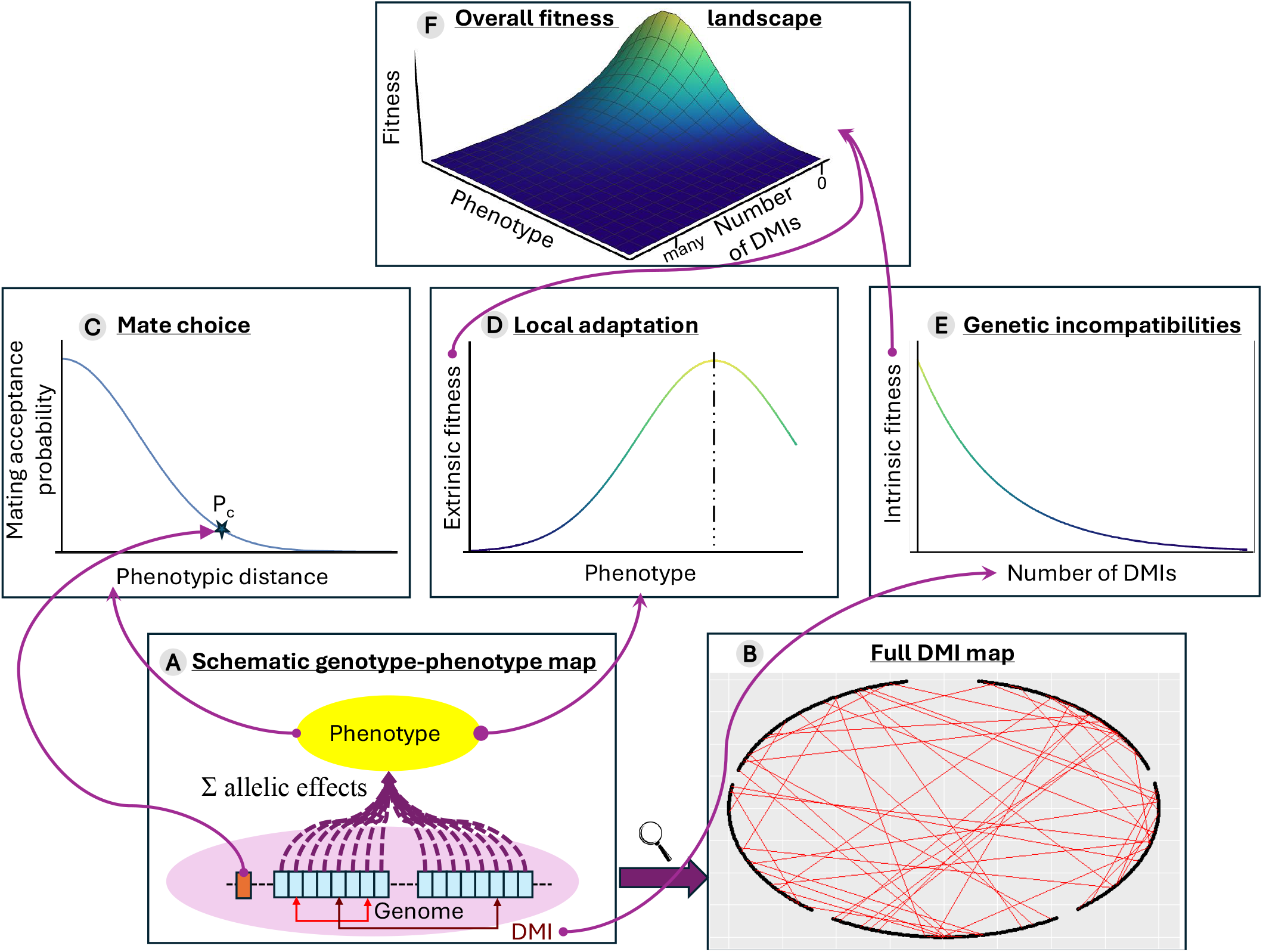
Schematic illustration of the genetic basis of the different mechanisms of reproductive isolation (RI). The diploid genome (panel A) consists of *L* loci separated in *n*_*LB*_ linkage blocks depicted by the blue squares. Their effect on each phenotypic trait are summed to generated the phenotype, depicted in yellow. In addition, a modifier locus (in dark orange) controls the strength of choosiness (*P*_*C*_), which determines how steep the probability of accepting a mating is (panel C, see eq. (3)). The phenotype determines the extrinsic fitness through a quadratic function (panel D, see eq. (1)). Finally, genetic incompatibilities are depicted by the red arrows between blue squares (panel B for the full map for the “default” scenario), and their number determined the intrinsic fitness (panel E, see eq. (2)). The overall fitness (before mate choice) is obtained by combining extrinsic and intrinsic fitness, as depicted in panel F.

The life cycle of individuals included the following sequence of events at each generation: (i) selection (based on intrinsic and extrinsic fitness components), (ii) migration, (iii) mating and (iv) zygote formation (including recombination and mutation). We considered two possible life cycles differing in whether selection or migration occurred first, i.e., differing in the dispersal stage of the organism. With “adult dispersal”, selection occurred before migration, whereas with “juvenile dispersal” selection occurred after migration. Since fitness of an individual was computed when it reached adulthood, the difference between the two life cycles was caused by the dispersal stage and whether immigrating individuals had their fitness determined by the environment they were born in (“adult dispersal”) or by the environment they inhabited at the time of reproduction (“juvenile dispersal”).

Migration was modeled as a Poisson process, of parameter *N×m* (per default, *m* = 0.05 and *N* = 10, 000). We investigated migration ranging from 5×10^−5^ to 0.2, corresponding to 2*Nm* values between 1 to 4,000, covering most of the reported values in Hey and Pinho (2012) (we did not considered lower values as *m* = 5 × 10^−5^ and *m* = 0 generated the same patterns in our simulations). Each population produced *N* offspring with a 50:50 sex ratio.

### Genotype and phenotype

The genome of each diploid individual was composed of two different (autosomal) elements: a modifier locus determining mate choice and one large genomic element determining the phenotype and genetic incompatibilities, composed of *L* biallelic loci (per default, *L* = 500) distributed equally over *n*_*LB*_ linkage blocks (fixed to *n*_*LB*_ = 5). These *L* biallelic loci determined the *n*_*t*_ quantitative traits (per default, *n*_*t*_ = 1). Mutation was bidirectional, symmetric and happened at rate *µ* (per default, *µ* = 10^−6^ per generation per locus). All mutations were pleiotropic and isotropic (affecting all *n*_*t*_ phenotypic traits equally), with phenotypic effects 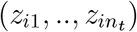 for the *n*_*t*_ traits drawn from a multivariate centered normal distribution with the covariance matrix 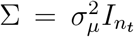, where 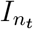 is the identity matrix and 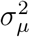 the variance of mutational effects (fixed to *σ*_*µ*_ = 0.05). Mutational effects were additive at the phenotypic level, hence the phenotype of a (diploid) individual was given by 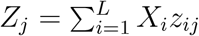, for a given phenotypic trait *j*, with *X*_*i*_ taking the value 0 if the individual was homozygous for ancestral allele, 1 if heterozygous and 2 if homozygous for the derived allele at locus *i*. For simplicity, we will refer to the phenotype that is at the optimum in the focal environment as the “optimal phenotype”, and to the phenotype at the optimum in the other environment as the “alternative phenotype”. Given that we only explored a low number of phenotypic traits (1 to 3), we did not correct the mutational effect size by the size of the phenotypic space. Recombination between adjacent loci (within a block) happened at rate *ρ* (fixed to *ρ* = 10^−3^ per individual and per generation), while recombination between linkage blocks happened at rate *ρ*_*LB*_ (fixed to *ρ*_*LB*_ = 10^−2^). For mate choice, we modeled choosiness as a single modifier locus (Haldane, 1941; Nei, 1967; Feldman et al., 1996) with multiple alleles, rather than a quantitative trait with many underlying loci.

Mutation happened at a fixed rate 10*µ* at the choosiness locus and the new allele differed from the current allelic value by a deviation drawn from a normal centered distribution with variance 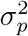 (fixed to *σ*_*p*_ = *σ*_*µ*_ = 0.05). The effect of alleles was additive and allelic values exceeding the boundaries (< 0 or > 0.5) were set to the corresponding boundary value. The choosiness locus was situated at the beginning of the main genomic element, at a genetic distance *ρ*_*MC*_ from the first position of the first linkage block (fixed to *ρ*_*MC*_ = 0.1).

Sex was not genetically determined, and was randomly assigned to each individual at birth according to the expected sex ratio (50:50).

### Fitness

The fitness *w* of an individual before mate choice was obtained by considering both extrinsic 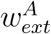 (due to the environment, here A) and intrinsic (due to the genetic incompatibilities) *w*_*int*_ components: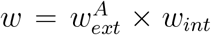, and corresponded to the relative probability of an individual to initiate a mating. Therefore, since *w* does not account for mate choice, it represents a component of fitness.

### Local adaptation (extrinsic fitness)

The extrinsic component of fitness was modeled using Fisher’s Geometric Model (Fisher, 1930), as it captures the distribution of fitness effect of mutations across different organisms (e.g., vesicular stomatitis virus, *Escherichia coli, Drosophila melanogaster* or *Triticum durum*, Martin and Lenormand, 2006) or *Arabidopsis thaliana* (Stearns and Fenster, 2016). The (absolute) fitness of each individual depended on the (Euclidean) distance between their phenotype, given by the 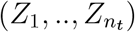 vector over the *n*_*t*_ phenotypic traits, and the environmental optimum. The relation between phenotype and fitness was obtained from Gros et al. (2009) with a slightly different choice of parametrization, to explicitly account for fitness in the alternative environment (see SI section M1). The fitness of an individual in environment A was given by:

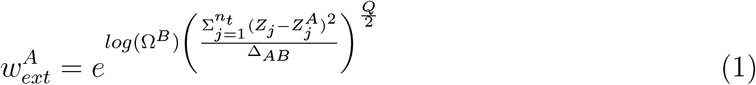

where *Z*_*j*_ is the phenotypic value of trait *j*, 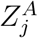 is the optimal phenotypic value for trait *j* in environment A, Δ_*AB*_ is a constant corresponding to the squared (Euclidean) distance between the phenotype of two individuals at the optima in environment A and environment B 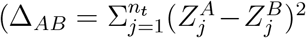 ; per default, Δ_*AB*_ = 100), *Q* controls the shape of the fitness landscape (*Q* ∈ [0, ∞[; per default, *Q* = 2), and Ω^*B*^ is a constant corresponding to the fitness of an individual in environment A with a phenotype perfectly adapted to environment B 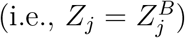. In this fitness landscape (Gros et al., 2009) when *Q* < 2, epistasis between pairs of mutations at the optimum was positive on average, otherwise it was negative (with null epistasis for *Q* = 2). The fitness in environment B 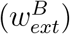 was obtained by substituting 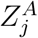 by 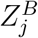, and Ω^*B*^ by Ω^*A*^ (per default, Ω^*A*^ = Ω^*B*^ = 0.1). The shape of the landscape determined the fitness of the F_1_ individuals with regards to the parents (see SI section M2 and equation (S3)). Under the default parameters (Table 1), weak hybrid vigor was the default behavior (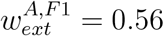 was larger than 0.55, the mean fitness of parents).

### Genetic incompatibilities (Intrinsic fitness)

We modeled intrinsic fitness, as pairs of Bateson-Dobzhansky-Muller incompatibilities (DMIs, Bateson, 1909; Dobzhansky, 1936; Muller, 1942). Importantly, as mentioned above, the loci involved in incompatibilities also affected the phenotype and therefore could be involved in local adaptation. We considered *n*_DMI_ pairs of incompatibilities (per default, *n*_DMI_ = 50), with the loci chosen among the *L* loci that affected the individual phenotype. Per default, each locus could form at most a single DMI, with the two interacting loci chosen independently of their position along the genome. Each DMI was codominant and their interaction multiplicative, resulting in the following formula for the intrinsic fitness component:

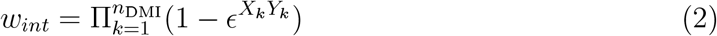

where *X*_*k*_ and *Y*_*j*_ are the counts of the number of derived alleles (0, 1 or 2) at the two loci forming the DMI, and *ϵ* is the strength of epistasis (i.e, the reduction in fitness due to a single DMI; per default, *ϵ* = 0.1). As extensions, we explored some alternative genetic architectures (e.g., DMIs between loci not affecting the phenotype), detailed in the Appendix.

### Mate choice

We modeled mate choice using a phenotype matching approach following Irwin (2020), where the probability of accepting a mate depended on the phenotypic distance between the two individuals, and was given by a Gaussian function. Rather than defining a variance, the shape of the Gaussian function was controlled by one parameter, *P*_*C*_ (Fig. S1), that corresponded to the probability of an individual with the “optimal phenotype” accepting a mate with the “alternative phenotype” (see SI section M3). This model could applied to either male or female choice as no aspect is sex-specific, and both choosing and chosen individuals can mate multiple times. Here, motivated by the spider mite example (Potter et al., 1976), we assumed that males were the choosy individuals. While mate choice has often been modeled as female choice (e.g., Servedio and Boughman, 2017) and male choice is rarer and possible under stricter conditions (Fitzpatrick and Servedio, 2018), it is still frequently found in nature (reviewed in Barry and Kokko, 2010). Mate choice proceeded as followed: a male and *χ* potential females (sampling with replacement) were chosen (fixed to *χ* = 100), according to their respective fitness. Second, the male considered mating with the first potential female, accepting said mating according to his choosiness (*P*_*C*_) and how phenotypically similar the two potential mates were, with probability

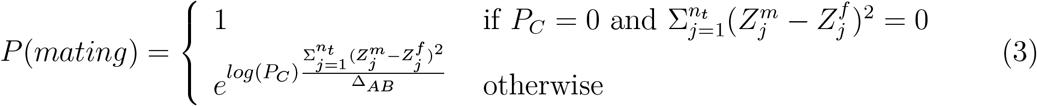

where 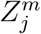 and 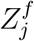 are the phenotypes at trait *j* of males and females, respectively. If the mating was accepted, a single offspring was produced and a new set of one male and *χ* females were selected. If the first potential female was rejected, the male would be considered for mating with the second female. This process continued until a mating was accepted or all *χ* females were rejected. If a male failed to accept any of the *χ* potential females, the male was discarded (but available to be chosen to initiate a mating again based on his fitness). This process continued until *N* offspring were produced. The number of potential females considered by a male for a mating (*χ*) therefore corresponds to a cost of choosiness, ranging from costly (*χ* = 1) to cost-free (*χ* → ∞). As mentioned above, choosiness (*P*_*C*_) was determined genetically by the modifier locus, with values varying between 0 (strict assortative mating) and 1 (no choosiness) at the phenotypic level, and therefore between 0 and 0.5 at the allelic level.

### Starting conditions and population metrics

We let RI barriers evolve for 50*N* generations, considering three possible starting conditions: an “Ancestral” state, a “Secondary contact” (SC) state and a “Secondary contact with initial mate choice” (SC + initial mate choice) (as detailed in the Appendix). Regardless, populations were always in a monomorphic state at the start of the simulations (i.e., all individuals were identical in each population).

We used the following genetic map as “default”. It consisted in *L* = 500 loci affecting a single phenotypic trait (*n*_*t*_ = 1). Among them, 100 loci formed *n*_DMI_ = 50 DMIs pairs, with each locus involved in at most a single DMI pair. This is the map illustrated in Figure 1B. A complete list of the genetic maps investigated are described in Appendix.

For each barrier, we used a specific metric to capture its evolution through time. For local adaptation, we measured the within population mean phenotype, for mate choice, the within population mean choosiness and for the accumulation of DMIs, we used two metrics: the between population intrinsic hybrid load and the number of equivalent fixed DMIs needed to generate the observed hybrid load (detailed in Appendix). Finally, we measured RI as the reduction in gene flow at a neutral marker, following Westram et al. (2022). We used the sojourn time of a neutral allele (number of generations until loss) introduced by transferring one individual from one population to the other. This was measured in isolated populations; further details are given in Appendix.

Simulations were written using C++ (2011), with the gsl (v2.7, Galassi et al. (2002)) and boost (v1.80) libraries. Analysis was conducted using Mathematica (v12, Wolfram Research, Inc. (2019)), RStudio (v2022.07.2+576) and R (v4.2.0, R Core Team (2022)), with figures generated using the ggplot2 package (v3.5.1, Wickham (2016)).

## Results

### Evolution of strict mate choice or accumulation of genetic incompatibilities depends on migration rate

We identified two possible regimes in the evolution of the three mechanisms of reproductive isolation (RI; Fig. 2). Local adaptation was always the first mechanism to establish (reaching 90% of the mean final value took between 0.2*N* and 0.3*N* generations for *m* = 0, and 1.4*N* to 11*N* for *m* = 0.2), but whether it was followed by the build-up of intrinsic postzygotic genetic barriers (accumulation of genetic incompatibilities) or the evolution of prezygotic barriers (strict conspecific mate choice) depended on the migration rate. For convenience, we referred to migration values leading to the first regime as weak and and the second as strong. Although the mean population phenotypic values moved further away from the environment optimum (here *z*^*A*^ = 5) as migration rate increased, migration did not prevent local adaptation, as a large proportion of individuals had phenotypes close to the optimum and similar to the ones of isolated individuals (Fig S2). We noticed that local adaptation stopped occurring in some replicates when migration became close to extrinsic hybrid load 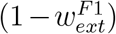 (Fig. S3). We did not find any case where the three barriers evolved simultaneously consistently across all 100 replicates (Fig. 3A). There were only 27 cases out of 890 where local adaptation, strict mate choice, and nonnegligible hybrid load evolved between populations (using the minimum mean hybrid load or choosiness observed in isolation as thresholds - i.e., hybrid load > 0.194 and choosiness < 0.034).

**Figure 2:**
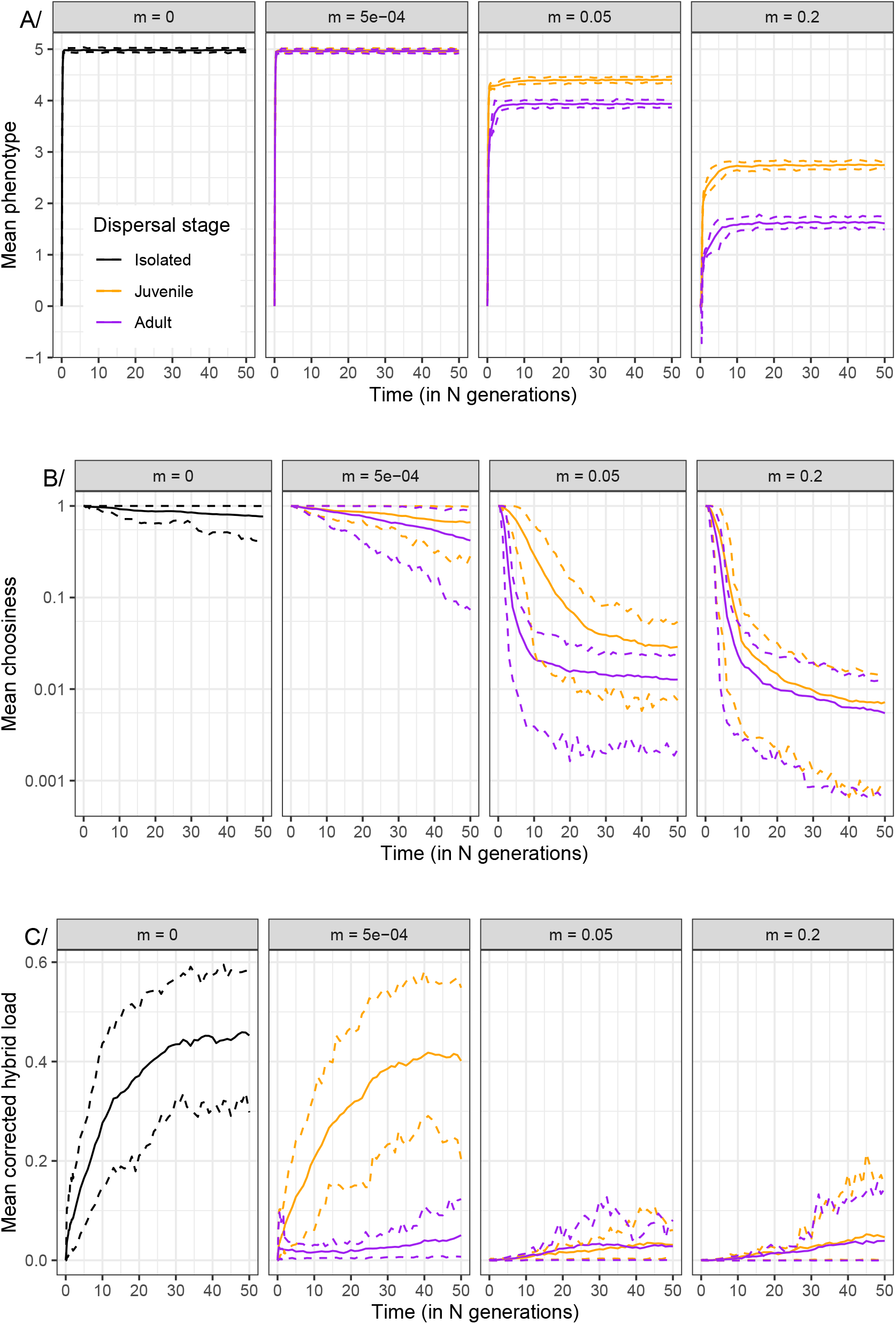
Evolution (in population A) of the phenotype (panel A), mate choice (B; on a log scale) and corrected hybrid load (C) for different migration rates (given in the header) and different dispersal stages over 50*N* generations. The solid lines correspond to the mean over the 100 replicates, and the dashed lines to the 10^*th*^ and 90^*th*^ percentiles. Relation between the evolution of mate choice and corrected hybrid load are displayed in Figures 3 and S7. Stronger RI barrier corresponded to convergence of mean phenotype to 5, mean choosiness to 0 and mean corrected hybrid load to 1. For reference, the migration rates given in the headers correspond to 2*Nm* values of 0, 10, 1000 and 4000, respectively. The genetic map used was the “default”, populations started in the “ancestral” state and other parameters correspond to the default values given in Table 1.

**Figure 3:**
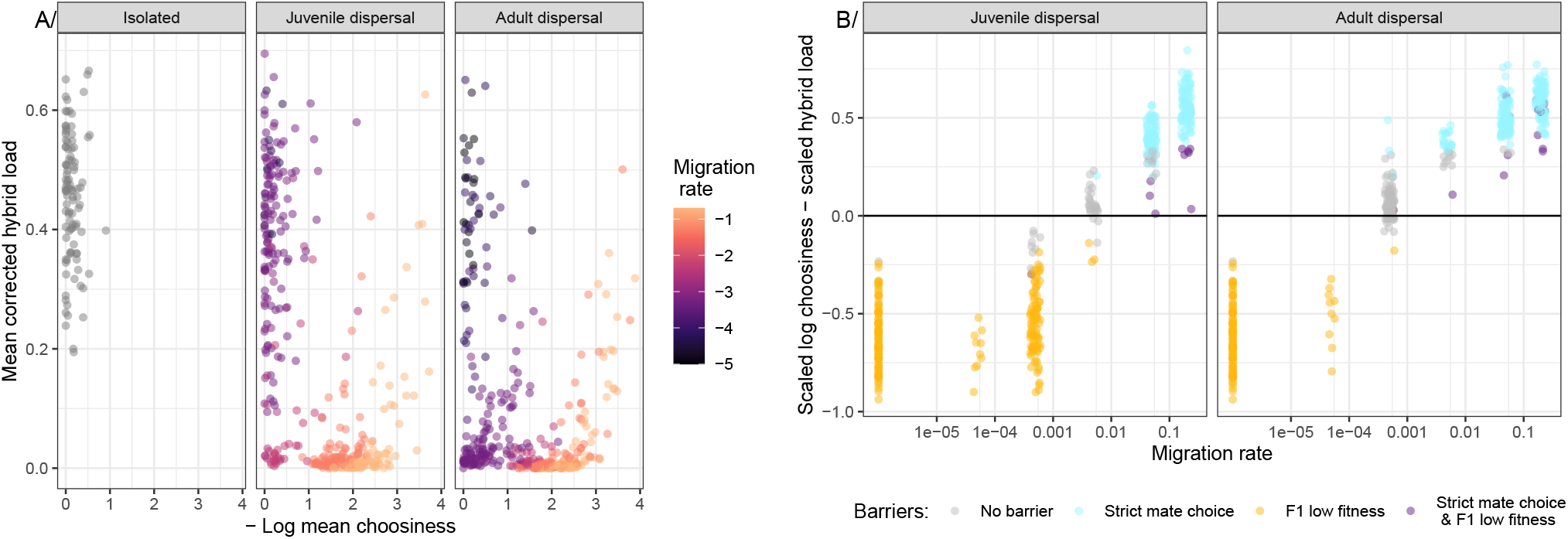
Evolution (in population A) of strict mate choice and accumulation of DMIs are (mostly) exclusive and depends on the migration rate. A/ Relation between (-log) choosiness (strict mate choice corresponds to large values) and mean corrected hybrid load (higher values indicate DMI accumulation) at the end of the 50*N* generations. Color indicates the migration rate (on a log-scale). A detailed figure of this relation per migration rate is given in Figure S7. B/ Relative contrast between scaled log choosiness (defined between 0 (no mate choice) and 1) and scaled hybrid load (defined between 0 (no hybrid load) and 1) as a function of migration rate (*m* + 10^−6^ on a log-scale) and dispersal stage. Color indicates whether RI mechanisms evolved (beyond local adaptation that always evolved). We defined evolution of the RI mechanisms using the minimum values observed in isolation; i.e., “No barrier” corresponded to a hybrid load ≤ 0.194 and choosiness > 0.034, “Strict mate choice” to a hybrid load ≤ 0.194 and choosiness ≤ 0.034, “F1 low fitness” a hybrid load > 0.194 and choosiness > 0.034 and “Strict mate choice & F1 low fitness” a hybrid load > 0.194 and choosiness ≤ 0.034). Other parameters correspond to the default values and are given in Table 1.

In the absence of migration, populations evolved toward their phenotypic optimum. In the weak migration regime, once local adaptation has evolved, it was followed by the accumulation of DMIs, but no evolution of mate choice. This regime of local adaptation, (almost) random mating and DMIs accumulation across populations extended to weak migration rates, with the threshold depending on the dispersal stage (up to *m* < 5×10^−5^ for “adult dispersal” and *m* < 5 × 10^−4^ for “juvenile dispersal”, Fig. 3, S4). Mate choice behaved neutrally in this context, but was displaced to values lower than *P*_*C*_ = 1 by mutational bias. Indeed, with weak migration, evolving towards strict assortative mating (*P*_*C*_ → 0) was not advantageous since the population was relatively homogeneous. However, there was no selective pressure to be less choosy  *P*_*C*_ → 1; i.e., random mating), as males were allowed to consider multiple females when reproducing (i.e., for *χ* > 2 the cost of being choosy was lower than the effect of drift, Fig. S5).

In the moderate to strong migration regime, once populations became locally adapted, strict mate choice evolved, but surprisingly, despite evolution of prezygotic barriers, there was almost no accumulation of DMIs. This corresponded to a regime of local adaptation, strict assortative mating and few or no DMIs. The evolution of strict mate choice (*P*_*C*_ < 0.034) can be explained by matings between individuals, adapted to the two environments, producing individuals with both unfit phenotypes (low extrinsic fitness component), and with potential expression of DMIs (low intrinsic fitness component). Thus, there was a strong selective pressure towards the evolution of assortative mating to avoid such unfit matings. Conversely, under these conditions, gene flow was still prevalent enough to prevent the accumulation of DMIs.

That is, we found that despite strong extrinsic and prezygotic barriers to gene flow, both in terms of local adaptation (the optimum phenotype of environment A was ∼ 10x fitter than an immigrant with the optimum phenotype of environment B, and 1.78x fitter than the F_1_ phenotype), and mate choice (a male at optimum phenotype of A will accept mating with a immigrant female with the optimum phenotype of B in ∼ 1% cases), in most cases the between-population hybrid load remained close to 0 when migration rate was higher than 0.005. We expected that incompatible alleles expressed in F_1_s would create an intrinsic hybrid load that became stronger with increasing migration rates, due to a higher proportion of F_1_ individuals. However, provided that there was sufficient gene flow (i.e., through F_2_s and further backcrosses), selection against incompatible alleles involved in DMIs should prevent their accumulation (seen for *m* ≥ 0.005). Alternatively, very strict mate choice could result in a low mean hybrid load if immigrants behave as an isolated population in the alternative environment. This was not the case, as the presence of gene flow was further confirmed by the multimodal distribution of phenotypes after 50*N* generations (Fig. S2), with peaks corresponding to F_1_ individuals (phenotypic value close to 0), and backcrosses with intermediate phenotypes (F_2_s around *±*2.5). The absence of incompatible alleles at DMI loci was further confirmed by checking the genomes of individuals (after 50*N* generations; Fig. S6).

The dispersal stage influenced the migration rate at which we saw a transition between the two regimes (Fig. 3B), with a lower migration rate threshold with “adult-dispersal”. This is expected because immigrating individuals with maladapted phenotypes were more likely to mate in the “adult dispersal” regime (their fitness depends on their birth environment), leading to stronger selective pressure to avoid them. Interestingly, we did not find a parameter space where both strict mate choice and DMI evolved simultaneously (and consistently) at significant strength (Fig. 3B, no values of *m* with a majority of purple points). As migration increases, two selective pressures, absent in isolation, emerged: selection for stricter mate choice and selection against DMIs. Our results suggested that selection against incompatibilities was more effective than selection for stricter mate choice, resulting in a Goldilocks zone, where neither strict mate choice evolved nor DMI accumulated.

Yet, within a given set of parameters, we observed a correlation between stricter mate choice and higher hybrid load, suggesting a trend for the two mechanisms to coevolve, in particular for moderate to strong migration rate (Fig. S7). This was likely a consequence of evolution of mate choice: once it became strict enough (*P*_*C*_ < 10^−3^), gene flow was sufficiently reduced such that the two populations behaved as isolated, and DMIs would start accumulating (Fig. 3, S8). As a consequence, we expected reinforcement (i.e., the evolution of strict mating due to genetic incompatibilities) to be unlikely under our model.

To further confirm the absence of reinforcement, we investigated secondary contact cases, with or without initial strict assortative mating. Under all considered scenarios, the postzygotic barrier due to DMIs eroded over time, either until it was mostly gone or towards some hybrid load equilibrium (Fig. 4). When secondary contact occurred between populations without initial mate choice (“SC” starting with *P*_*C*_ = 1), we found that the evolution of mean choosiness towards strict mate choice was similar to when two populations diverged from an ancestral population (“Ancestral”), and the hybrid load due to DMIs converged to similarly low values (mean hybrid load ≤ 0.10) in both scenarios (Fig. 4 left vs mid panels). In contrast, when secondary contact occurred between populations already with DMIs and strict mate choice (“SC + initial mate choice” starting with *P*_*C*_ = 10^−4^), we found a distinct dynamics, with maintenance of very strict mate choice for scenarios with strong migration (*m* > 0.05), and evolution of hybrid load towards high equilibrium values (mean hybrid load ∼ 0.35), irrespective of the migration rate. Given that the erosion of the postzygotic barrier due to DMIs happened relatively slower than the evolution of choosiness, this supports effective purging of DMIs.

**Figure 4:**
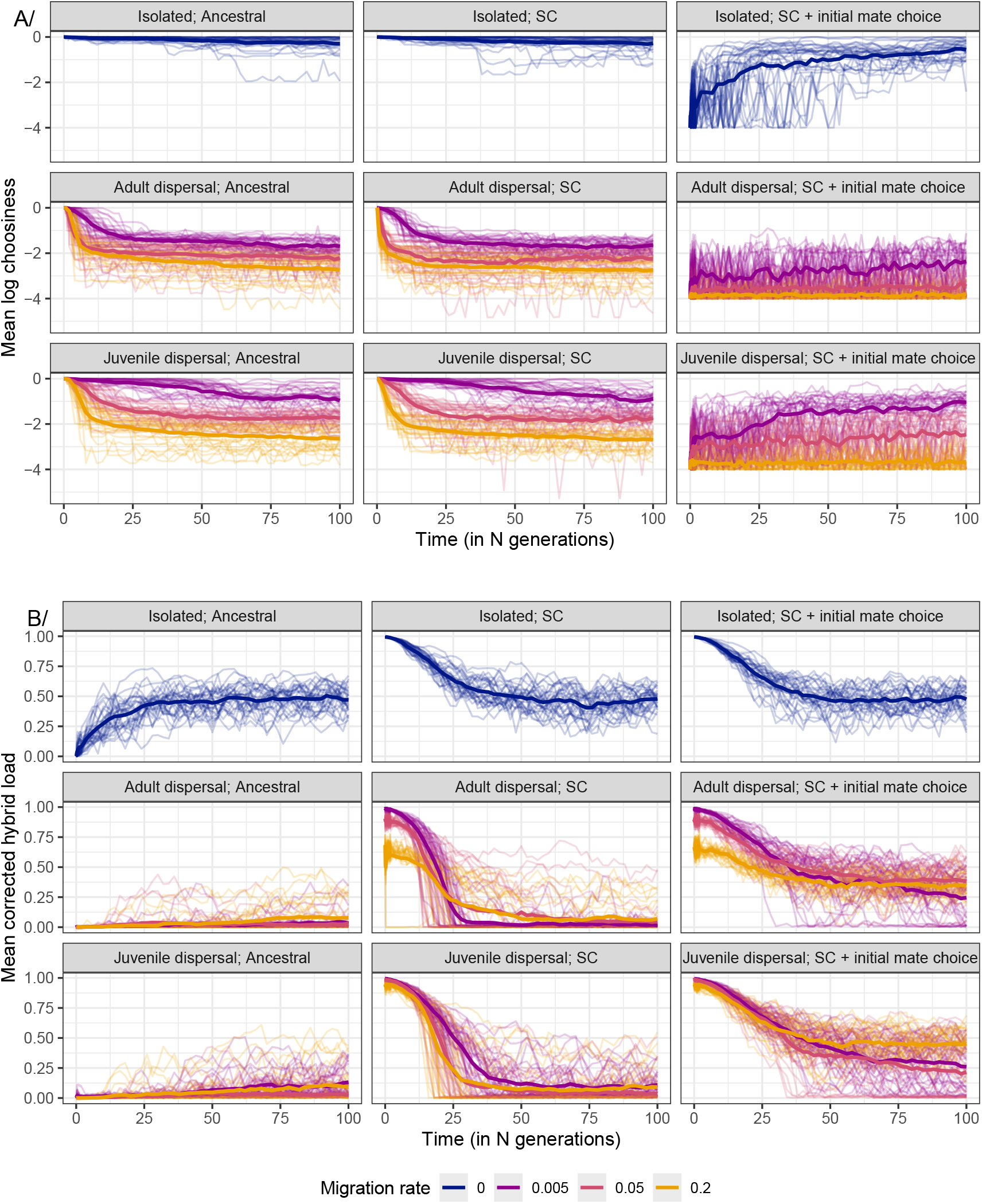
Evolution (in population A and over 100*N* generations) of mate choice (panel A) and corrected hybrid load (B) for different dispersal stages and different starting conditions: the “Ancestral” (default) scenario, the secondary contact (“SC”) scenario and the secondary contact with initial mate choice scenario (“SC + initial mate choice”). Different migration rate are given indicated by different colors, with each thin line corresponding to a different replicate and the thick line to the mean over the 30 replicates. The final states and link between mate choice and hybrid load are displayed in Figure S9. Other parameters correspond to the default values and are given in Table 1

Overall, this indicates that reinforcement (as defined above) is not possible under the investigated scenarios, due to the effective removal of DMIs when there is enough gene flow (*m* ≥ 0.005), consistent with previous theoretical work on continent-island models (Agrawal et al., 2011; Bank et al., 2012).

We considered alternative genetic architectures to better understand how the shared genetic architecture between local adaptation and a second RI mechanism shaped their interactions. First, in the absence of local adaptation (Fig. S10), mate choice and DMIs alone could not maintain the population differentiation, either due to a collapse of the RI barrier (at weak migration rates) or ecological exclusion of one of the populations (at strong migration rates, SI section R2). Second, having the DMI loci involved in local adaptation (as presented above) or not (“neutral map”) led to a similar evolution of each of the three RI barrier investigated (Fig. 5, SI section R4). Finally, when mate choice was based upon assortative mating at a neutral phenotypic trait, mate choice was unable to evolve from an ancestral state nor from a secondary scenario (Fig. S11), but could persist if already present, and if F_1_s intrinsic fitness was sufficiently low (SI section R3).

**Figure 5:**
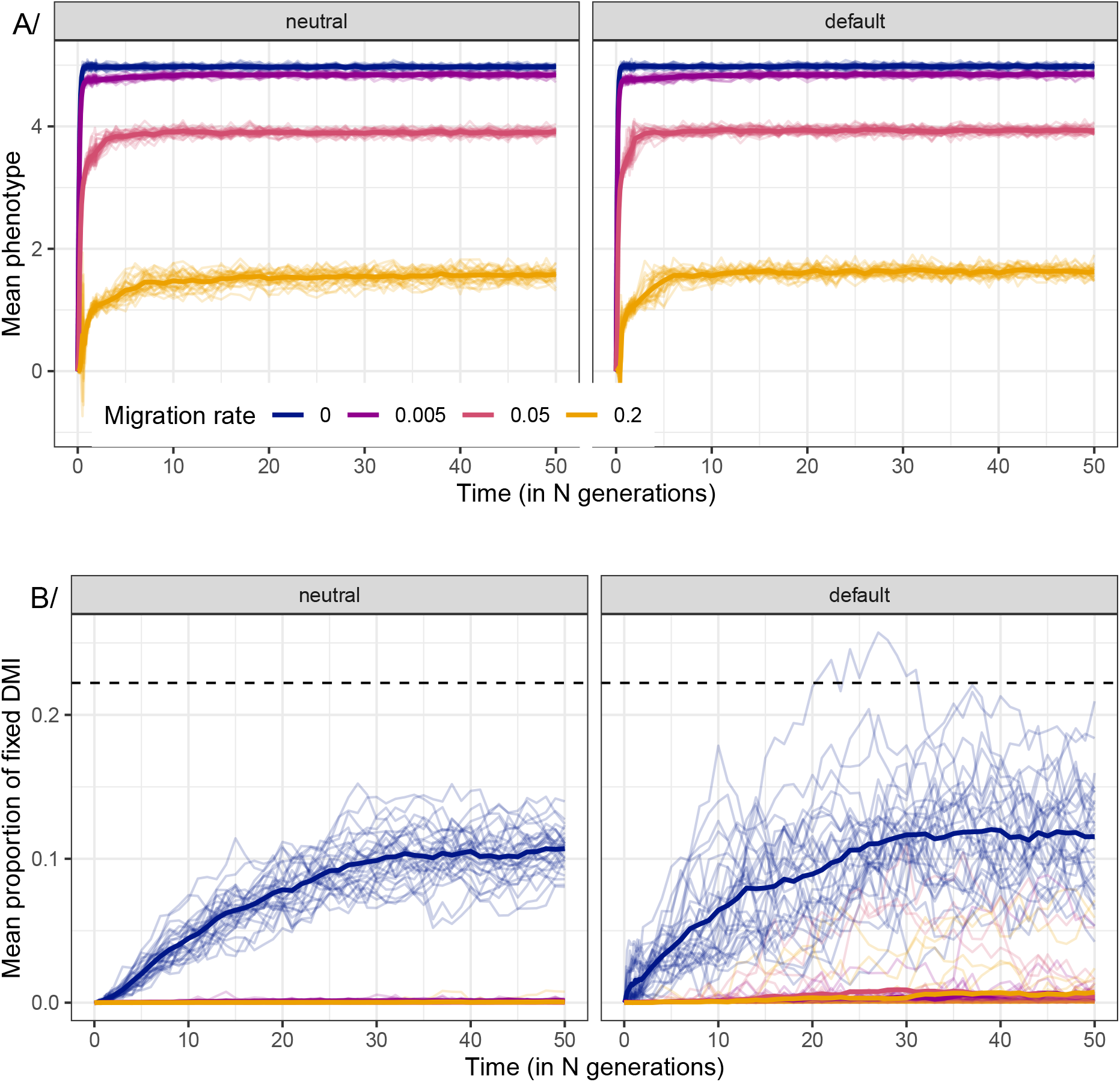
Evolution (in population A and over 50*N* generations) of the phenotype (panel A) and number of equivalent fixed DMIs (B; computed according to equation (A1)) for different DMIs architecture (given in the header) and different migration rates. Each thin line corresponds to a different replicate, with the thick line corresponding to the mean over the 30 replicates. Color corresponds to different migration rates: *m* = 0 in blue, *m* = 0.005 in purple, *m* = 0.05 in pink and *m* = 0.2 in orange. For the B panel, the black dashed line corresponds to the weak mutation strong selection approximation for the proportion of DMI fixed between populations, 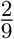. For the “neutral” architecture, DMIs are formed between loci not affecting the phenotype, while for the “default” architecture, the DMIs were formed between loci affecting the phenotype. Other parameters correspond to the default values and are given in Table 1

### No accumulation of DMIs with migration irrespective of the network of DMI interactions

Across migration rates, the intrinsic hybrid load was either absent or evolved towards some non-zero equilibrium value (Figure 2C). In the later case, the hybrid load matched what was observed in the absence of gene flow (*m* = 0), and could be obtained starting from an ancestral population or a secondary contact scenario. Such equilibrium hybrid load was equivalent to having five DMI pairs differently fixed between the two populations. This indicated that in scenarios where DMIs did accumulate, only about 10-15% of all possible DMIs were differently fixed between the two populations (in terms of equivalent number of DMIs; Fig. S12). This could be partially explained by the design of the genetic architecture and mutation process, as forward and backward mutation rates (at each biallelic locus) were considered equal.

To understand why only 10 to 15% of the DMIs were differentially fixed, we considered a simple deterministic model describing how DMIs would accumulate between populations in the limit of weak mutation and strong selection (i.e., strong epistasis, *ϵ* → ∞). In each population, at equilibrium and assuming a low mutation rate, about half of the loci would be fixed for the derived allele (i.e., *A*_*i*_ or *B*_*i*_ at loci A and B of DMI pair *i*) and the other half for the ancestral allele (i.e., *a*_*i*_ or *b*_*i*_). In addition, for each DMI pair, the derived alleles (*A*_*i*_ or *B*_*i*_) could only arise in the ancestral background, whereas the ancestral alleles could arise in any background. Under these conditions, the population had three possible states: fixed for *a*_*i*_*b*_*i*_, *A*_*i*_*b*_*i*_ or *a*_*i*_*B*_*i*_. The expected probability of having *A*_*i*_ fixed in the first population at a given time-point was 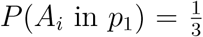 (see SI section M5 for details). Therefore, the probability of having a DMI between the two isolated populations was 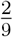, while in the absence of back mutation, this probability rose from 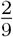 to 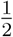.

We observed a lower number of DMIs in the simulations than predicted by the weak mutation strong selection approximation. This was the case irrespective of the epistasis strength (*ϵ* = 0.9 or *ϵ* = 0.01, Fig. S13), or the genetic map (“default”, “neutral”,”network” or “optimized” maps). Yet, we found that the scaled mutation rate (i.e., *θ* = 4*Nµ*) influenced the proportion of equivalently fixed DMIs (Fig. S14). These results suggested that, in large populations or when mutation rate is high, multiple mutations might segregate at the same time within each population. If both *A*_*i*_ and *B*_*i*_ were polymorphic simultaneously in a given population, selection against those incompatible alleles would prevent their fixation. Indeed, in isolated populations at generation 50*N*, almost 40% of all pairs of DMIs had both alleles *A*_*i*_ and *B*_*i*_ segregating in the population, yet with the rarer among the two alleles never exceeding a frequency of 0.015 (Table S1).

### Local adaptation as the main barrier to gene flow in scenarios with migration

To quantify the barriers to gene flow we computed the mean sojourn times of a neutral allele from a focal immigrant individual. Regardless of the migration rate at which populations evolved, the mean sojourn time of an unlinked neutral marker was usually lower or similar to the expected sojourn times under neutrality, indicating that some RI barriers effectively reduced gene flow (e.g., local adaptation) where other did not or did not evolve (e.g., mate choice at weak migration rates, and genetic incompatibilities at strong migration rates, Fig. 6). The combination of barriers always decreased the mean sojourn time compared to the effect of each barrier individually. This result was qualitatively the same if we considered a linked neutral marker instead (Fig. S15). This was the case regardless of the sex of the focal individual, although the choosy sex (males) had longer sojourn times when mate choice was involved (“MC”, “LC+MC”, “MC+DMI”, “all”), consistent with higher gene flow when immigrants were of the choosy sex. Indeed, we demonstrated that as long as choosiness was not too costly, the choosy sex was responsible for most of the gene flow between populations (see Appendix and Fig. A1).

**Figure 6:**
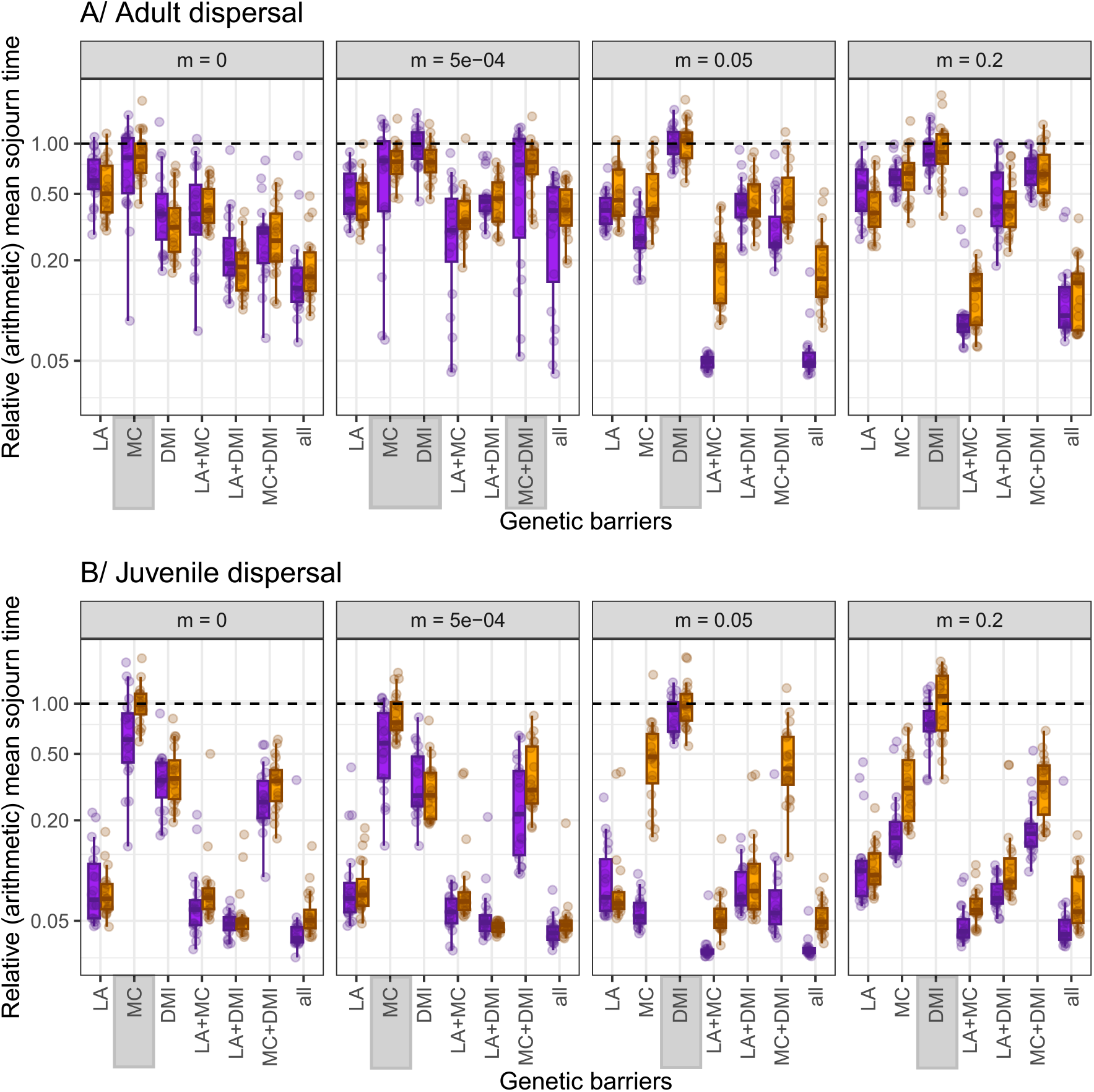
Impact of reproductive isolation (RI) barriers on the relative sojourn time of neutral alleles at a freely recombining marker introduced by a focal immigrant individual. The sojourn time corresponds to the number of generations until the introduced marker was lost and used as a proxy for the barrier strength, with lower values indicating stronger barriers against gene flow. Sojourn times for specific RI barrier were obtained by turning off the other RI barriers, and the x-axis indicates which barriers were kept active: local adaptation (LA), mate choice (MC), postzygotic isolation (DMI), a combination of two (e.g., LA+MC) or all three barriers together (all). Depending on the migration rate, some RI barrier may not have evolved; in these instances the RI barrier was highlighted in gray. Results shown for a male (orange) and female (purple) focal immigrant individual. Sojourn times were measured relative to the mean sojourn time under neutrality across all replicates, i.e., neutrality corresponds to a value of 1 (dashed horizontal line). Lower values indicate less introgression, i.e., stronger barriers. Panel A corresponds to the “adult dispersal” stage and panel B to the “juvenile dispersal” one. Additional migration rates are displayed in Fig. S16. The corresponding figures for the absolute times (Fig. S17), and for the relative sojourn time of a linked marker (Fig. S15) are given in Supplement. Finally, Fig. S19 displays the relationship between individual barrier strength and sojourn time. Other parameters correspond to the default values.

Across all combinations of barriers and dispersal stages, the shortest sojourn times were similar irrespective of the migration rate, indicating that RI barriers evolved to similar levels with and without migration. Irrespective of the dispersal stage and migration rates, cases where all barriers were active (“all”) showed shorter sojourn times, i.e., stronger barriers. Yet, cases with local adaptation (“LA”), local adaptation and mate choice (“LA+MC”) and local adaptation and genetic incompatibilities (“LA+DMI”) often led to similar sojourn times to those found by combining all barriers (“all”), although this depended on the migration rates and dispersal stage. In agreement with purging of DMIs, when migration *m* ≥ 0.05, the sojourn times when only the DMI barrier was active (“DMI”) were similar to neutral. In contrast, without migration, DMIs evolved successfully and acted as a RI barrier reducing sojourn times to ≈ 0.40 of the values observed under neutrality.

Interestingly, across most scenarios we found differences depending on the dispersal stage in the reduction of gene flow (Fig. 6). First, for scenarios with local adaptation (“LA”, “LA+MC”, “LA+DMI”, “all”), the reduction in sojourn time was stronger for “juvenile dispersal” than “adult dispersal”, i.e., local adaptation was a much stronger barrier to gene flow with “juvenile dispersal”. Second, for “juvenile-dispersal”, when all mechanisms were active (“all”), the reduction in sojourn time was mostly independent of the migration rate and of the sex of the focal immigrant individual, since focal individuals failed to reproduce in the first generation. For “adult-dispersal”, this was no longer the case, as sojourn times depended on migration rate and sex of focal individual (strongest joint effect for “all” barriers observed for *m* = 0.005, Fig. S4). The evolution of stronger barriers at intermediate migration rates with “adult-dispersal” was due to a balance between two mechanisms: increasing migration triggered the evolution of mate choice, but also ensured that at each generation there was a proportion of individuals with the alternative phenotype in the resident population. As migration further increased, this proportion individuals with the alternative phenotype became sufficiently large, that the focal immigrant was likely to successfully reproduce. Therefore, under the “adult dispersal” life cycle, we predict that strong barriers to gene flow leading to speciation would only evolve for scenarios with the “right” amount of migration.

A high variance (overdispersion) for the mean sojourn time under some scenarios was observed, mainly for barriers involving mate choice (“MC”, “LA+MC”, “MC+DMI”, “all”) and for intermediate migration rates (*m* = 0.0005 and *m* = 0.05). This overdispersion reflects the stochasticity across replicates, since in some replicates, populations evolved strict mate choice, while not in others.

Finally, we investigated the sojourn time of neutral alleles in scenarios of secondary contact with preexisting choosiness and fixed levels of the strength of each barrier, thus removing the stochastic variation in evolution of barriers across runs. With “juvenile dispersal”, local adaption remained the strongest barrier to gene flow, i.e., shorter sojourn times for “LA” (Fig. S18), with addition of mate choice (“LA+MC”) further reducing the sojourn times (but only in non-choosy sex, and when each barrier was relatively weak). Genetic incompatibilities (“DMI”) were the weakest of all barriers, providing shorter reductions in sojourn time than the other mechanisms, but equivalent to mate choice (“MC”) in the choosy sex. With “adult dispersal”, local adaptation (“LA”) was one of the weakest barriers. Surprisingly, although with “juvenile dispersal” genetic incompatibilities (“DMI”) were one of the weakest barriers, with “adult dispersal” “DMI” provided the strongest barrier (i.e., shorter sojourn times), even stronger than local adaptation and without sex-specific effects.

## Discussion

We investigated how the interaction of different mechanisms of reproductive isolation (RI) impacts divergence and gene flow between populations, under a polygenic model of local adaptation and genetic incompatibilities with a shared genetic architecture. For the evolution of RI, Table 2 provides a summary of general trends observed. In scenarios with migration, local adaptation was unsurprisingly always the first barrier to evolve, with either accumulation of genetic incompatibilities (DMIs) or the evolution of strict assortative mating evolving afterwards. The evolution of the last two barriers were somewhat mutually exclusive: whether strict mate choice or accumulation of DMIs evolved was dictated by migration rate, and to a lesser extent by the life cycle. Surprisingly, even in cases where local adaptation and strict mate choice evolved, DMIs acted as a very weak barrier, consistent with purging of genetic incompatibilities provided there was sufficient gene flow. Thus, reduction in gene flow (strength of RI) was mostly a consequence of the action of local adaptation, with mate choice being also effective for cases with sufficient migration. Importantly, we modeled here mate choice as male choice but the model is fully symmetric and all genetic elements were autosomal. Therefore, all conclusions apply to both male and female mate choice.

**Table 2:** Summary table of our results, indicating the general trend, within the parameter ranges explored. Each row corresponds to a different combination of genetic architecture, different initial conditions, and (possibly other parameter changes). We summarized the general trend in the evolution of phenotypic divergence (in most cases a proxy for local adaptation), evolution of strict mate choice, and accumulation of DMIs. Figures supporting the results are indicated below each statement, and not indicated when results were not shown.

| Genetic architecture | Initial conditions | Phenotypic divergence | Evolution of strict mate choice | Accumulation of DMIs |
| --- | --- | --- | --- | --- |
| Default | Ancestral | Yes (approx. up to $m=1-w(F_1)$)<br>Fig 2; Fig S4; Fig S4 | If migration is strong enough<br>Fig 2; Fig S4 | If migration is weak or very strict mate choice has evolved<br>Fig 2; Fig 3; Fig S4 |
|  | Secondary contact | Yes | If migration is strong enough<br>Fig 4A | If migration is weak or very strict mate choice has evolved<br>Fig4A |
|  | Secondary contact with initial mate choice | Yes | Remained (if no isolation)<br>Fig 4B | Maintained at a mutation selection equilibrium if mate choice remains sufficiently strict<br>Fig 4B |
| Default - no LA | Secondary contact with initial mate choice | No; Ecological exclusion for $m \geq 0.005$ or for $m \leq 0.0005$ converge to ancestral state due to mutational bias<br>Fig S10; Fig S21 | No; Evolving toward choosiness values above 0.1<br>Fig S10; Fig S21 | No; Lost either due to ecological exclusion for $m \geq 0.005$ or purged by selection if $0 < m \leq 0.0005$<br>Fig S10; Fig S21 |
| Neutral display | Ancestral | Yes at the selected trait; No at the cue trait<br>Fig S11 | No<br>Fig S11 | Only if m is really low<br>Fig S11 |
|  | Secondary contact | Yes<br>Fig S11 | No<br>Fig S11 | Only if m is really low<br>Fig S11 |
|  | Secondary contact with initial mate choice | Yes<br>Fig S11; Fig S22 | Maintained if epistasis caused by DMI is strong enough - loss of MC triggered by the loss of DMI<br>Fig S11; Fig S22 | Maintained at a mutation selection equilibrium if epistasis caused by DMI is strong enough<br>Fig S11; Fig S22 |
| Neutral | Ancestral | Similar to default case<br>Fig 5 | Similar to default case | Similar to default case<br>Fig 5 |
| Network | Ancestral | Similar to default case<br>Fig S20 | Similar to default case | No<br>Fig S20 |
| Optimized | Ancestral | Similar to default case<br>Fig S20 | Similar to default case | Similar to default case<br>Fig S20 |
| Default 3D | Ancestral | Same as default case if corrected for the distance between optimum<br>Fig S23 | Similar to default case<br>Fig S23 | Similar to default case<br>Fig S23 |
| Default - effect of Q | Ancestral | Overall similar to default case, but with weaker divergence as Q increases<br>Fig S24 | Same as default case<br>Fig S24 | Overall similar to default case, but with lower hybrid load as Q increases<br>Fig S24 |
| Default 3D - asymmetry of optimums | Ancestral | Similar to default 3D case<br>Fig S25 | Similar to default 3D case<br>Fig S25 | Similar to default 3D case<br>Fig S25 |

### Local adaption at the center of speciation with gene flow

Under our model with polygenic architectures, local adaptation played a major role in the evolution of RI as it was always the first barrier to evolve. The key role of local adaptation, in models of speciation with gene flow, had already been pointed out previously using two-locus models (Agrawal et al., 2011; Bank et al., 2012). In addition to being the first RI barrier to evolve, with “juvenile dispersal”, local adaptation was the main mechanism responsible for the reduction of sojourn times of neutral introgressed alleles, a proxy for the level of gene flow between populations. The importance of ecological speciation has received a lot of attention (as detailed in Nosil, 2012 - but see Anderson and Weir, 2022). Our results are in agreement with the role of local adaptation in ecological and speciation scenarios with gene flow, but bring an important nuance. Indeed, when considering the “adult dispersal” life cycle, local adaptation drives the evolution of phenotypic differentiation between the two populations, and mate choice or accumulation of DMIs only evolve once populations become locally adapted. However, provided that populations evolved strict mate choice or accumulated DMIs, the strength of these two barriers becomes similar to local adaptation, as we found for secondary contact models from fixed population states (Fig. S18). For both the evolution from an ancestral population or from secondary contact from fixed population states, the relative strength of local adaptation depended on the life cycle, being stronger with “juvenile dispersal”. Since the only difference between the two life cycles was whether the fitness of an immigrant individual was affected or not by the new environment, this indicates that local adaptation is mostly effective upon arrival of the immigrant individual, but becomes ineffective to reduce gene flow when the focal individual successfully reproduces. This result highlights the brittle nature of local adaptation/ecological differentiation as a genetic barrier to gene flow (Blanckaert et al., 2020).

### Evolution of mate choice when there is sufficient migration

In our model, mate choice was never able to function as an independent barrier. It evolved only in the presence of local adaptation and non negligible migration. Therefore it always evolved as a consequence of another barrier, and corresponds to a form of coupling of multiple RI barriers (sensu Butlin and Smadja, 2018).This delay in the evolution of mate choice versus local adaptation has been previously reported across different mate choice models (Thibert-Plante and Gavrilets, 2013) when investigating prezygotic barriers exclusively.

Mate choice was also effective at limiting gene flow between species, but only for the non-choosy sex and when a very strict choosiness evolved. This is in line with previous simulations studies, highlighting the relatively ineffective role of assortative mating to maintain a narrow hybrid zone between populations (Irwin, 2020). As in Irwin (2020), our results partly stem from modeling choosiness using a Gaussian function and a matching phenotype approach. In particular, this implies that mating of F_1_s with identical phenotypes is very likely, and F_1_s are also likely to be accepted as potential partners by resident individuals. Our approach to model mate choice creates a “now or never” dichotomy for reducing gene flow, which is seen in differences between the choosy and non-choosy sex. If the focal immigrating individual manages to reproduce, then most of the RI barrier is gone and the introgressed marker behave somewhat similarly to a neutral mutation. Having the choosiness function more akin to a step function instead of a Gaussian could mitigate the effects described here and generate stronger barriers to gene flow. The suitability of each approach depends on whether the trait used for mate choice is qualitative or quantitative. In the later case, a continuous smooth function is a more likely representation of the biological process involved in mate choice (especially since we are not considering the environmental variance that could contribute to the phenotype and would flatten the choosiness function). Importantly, we never observed reinforcement, (i.e., the evolution of mate choice due to the presence of genetic incompatibilities), as the two mechanisms seem to exist in different parameter spaces. This is likely because the selection against DMIs, caused by the hybrid load, is more sensitive to migration than selection for stricter mate choice, resulting in a disjoint parameter space: combination of parameters where DMIs could accumulate did not overlap with the ones where stricter mate choice could evolve. While surprising given that occurrence of the two RI barriers is common in nature, we observed rare cases where both strict mate choice evolved and DMIs accumulated. In those rare cases, DMIs accumulated because of the evolution of mate choice, rather than the opposite. In addition, mate choice was not able to evolve if the trait used for assortative matings was not under local adaptation (i.e., if it was not a magic trait), but could persist if F_1_s individuals were sufficiently unfit. Finally, we chose to model mate choice as assortative mating, with both males and females being able to engage in multiple matings. This makes the results obtained here valid for both males and females.

### Genetic incompatibilities do not accumulate in locally adapted populations with strict mate choice when there is sufficient gene flow

Two types of incompatibilities were modeled in our approach: extrinsic and intrinsic ones. Indeed, Fisher’s Geometric Model naturally generates epistasis between mutations due to the quadratic relationship between phenotype and fitness. Within this framework, Fraïsse et al. (2016) showed that many empirical patterns of speciation (e.g., Haldane rule, heterosis, hybrid breakdown) can be mimicked, making it a useful model to study RI. However, such approach makes it extremely challenging to characterize the effect of genetic incompatibilities, as they depend both on the environment and the genetic background. Therefore, our model also included intrinsic incompatibilities using the classical Bateson-Dobzhansky-Muller model (Bateson, 1909; Dobzhansky, 1936; Muller, 1942). Assuming that the same locus affecting local adaptation can generate genetic incompatibilities may seem to be a strong assumption, yet examples of local adaptation generating DMIs have been reported (Macnair and Christie, 1983; Świadek et al., 2017; Frayer et al., 2025). In particular, adaptation of *Saccharomyces cerevisae* to low glucose medium (Anderson et al., 2010; Kvitek and Sherlock, 2011) or nyastin (Ono et al., 2017) reveals the existence of genetic incompatibilities between beneficial mutations. The same pattern was also observed in *Methylobacterium extorquens* (Chou et al., 2014). Regardless, we obtained the same results when the DMIs affected neutral loci, or when the architecture was “optimized” to favor DMI accumulation (Fig. S20), suggesting that local adaptation did not favor their accumulation.

Generally, under our model, genetic incompatibilities seem to arise more as a consequence of isolation (either due to weak migration rates or strong reproductive barriers), and not as an active mechanism to prevent gene flow, matching the predictions from a two-locus model (Bank et al., 2012). Our results also matched with the outcome of experimental evolution in Saccharomyces cerevisiae, where Dettman et al. (2007) observed the evolution of DMIs as a by-product of divergent selection between isolated populations. The same pattern was also observed in experimental evolution of isolated populations of *Neuspora* (Dettman et al., 2008). Whether such accumulation of DMIs as a by-product could happen with ongoing gene flow remain largely unexplored (Kulmuni and Westram, 2017). Our study reveals a duality in the effect of these intrinsic genetic incompatibilities in the speciation process. Accumulation of DMIs is relatively constrained, only happening when gene flow was absent or limited. This effect was further amplified when forward and backward mutation rates were symmetric, as the within-population mutational load generated by the possible DMIs tend to select against the fixation of any of the incompatible alleles (Fig S14), even in the absence of migration.

Despite this, multiple DMIs have been reported across many organisms (Presgraves, 2003; Kao et al., 2010; Corbett-Detig et al., 2013; Powell et al., 2020; Frayer et al., 2025), including a characterization of the underlying genetic mechanisms (reviewed in Table 1 of Kitano and Okude, 2024). Potentially, all of these DMIs may have formed in isolation, as described in the conceptual definition of the DMI model (Coyne and Orr, 2004) whether due to temporal, geographical or prezygotic barriers. However, isolation is rather unlikely, as gene flow has been reported for some of the cases mentioned above (e.g., between *Xiphophorus malinche* and *X. birchmanni* Powell et al., 2020). Alternatively, it is possible that DMIs accumulation happens at a much slower rate, as discussed for cichlids (Stelkens et al., 2010), that was not captured with our design here. Finally, DMIs are often thought as fixed differences between species. Yet as argued by Cutter (2012) and demonstrated in *Drosophila melanogaster* Corbett-Detig et al. (2013), this may not be the case, and many genetic incompatibilities may be segregating within populations - a feature that we recovered in our simulations (Table S1).

Under our model, as soon as migration was non negligible, genetic incompatibilities were unlikely to accumulate between populations, despite the evolution of phenotypic differentiation and strict mate choice. Interestingly, this pattern has been observed in nature in cases of ecological divergence. For example, cichlid fish species maintain their integrity despite coexisting with gene flow in at least parapatry, and in the presence of mate choice, yet hybrid breakdown remained limited (Stelkens et al., 2015) and DMIs accumulation happened at a much slower rate than speciation (Stelkens et al., 2010). Such pattern also matches the example of *Anopheles gambiae* and *A. coluzzi*. As summarized in Mallet and Mullen (2022), the two species display ecological differences and strong assortative mating, but are sympatric and apart from a few islands of differentiation, 99% of the genome is homogenized, and therefore not propitious to the accumulation of DMIs (Turner et al., 2005; Lee et al., 2013). The lack of characterized DMIs may be partly due to the challenge associated with their detection (Nouhaud et al., 2020; Blanckaert and Payseur, 2021), but our model here also highlights that low levels of gene flow between populations are sufficient to purge DMIs.

Finally, it is possible that some of the assumption of our model may limit the accumulation of DMIs. First, we are assuming a constant population size and therefore soft selection (Wallace, 1968, 1975) - low fitness phenotypes therefore have a higher chance of leaving offspring when frequent enough compared to a hard selection scenario. While extinction of either or both populations is a non negligible risk, demographics and evolutionary mechanisms have been shown to act in opposite direction with regards to the hybrid speciation process (Blanckaert et al., 2023). Therefore, DMIs may play a more important role when considering hard selection with both demographic and evolutionary mechanisms - such as allowing the populations to be affected by changes in effective population size. Second, mutations were limited to point mutations and structural variants (e.g., inversions, translocation) were not included. These large-effect mutations can be central to local adaptation and ecotype formation (e.g., 22 and 12 putative inversion have respectively been reported in *Littorina saxatalis* and L. fabalis (Johannesson et al., 2024), at least 4 putative inversions in *Coelopa frigida* (Mérot et al., 2021), and 3 inversions in Heliconius numata (Jay et al., 2021)), especially in the early stages of speciation (Kirkpatrick and Barton, 2006). Third, we focused on pairwise epistasis. However, higher order epistasis (Ayala-López and Bank, 2025) may play an important role, as it can partially alleviated the within-population mutational load, making the accumulation of DMIs more likely.

## Conclusions

We explicitly modeled the interaction between local adaptation, genetic incompatibilities (DMIs) and mate choice, by considering that the same loci could be involved in the phenotype (under selection in the two environments) and associated with DMIs, and that mate choice depended on phenotype and that choosiness was due to a modifier locus linked to sites determining the phenotype. Surprising, despite a common genetic basis that we expected to favor the joint evolution of all three RI barriers, we found that long term evolution from an ancestral population did not lead to this joint build up, i.e., locally adapted populations with strict mate choice and accumulation of DMIs. Instead, we found that populations evolved towards two regimes: either local adaptation and accumulation of DMIs with weak mate choice, or local adaptation with strict mate choice and no DMIs. This suggests two possible paths of speciation, one where ecological and genetic isolation dominate, and another where ecological and behavioral isolation dominate. The migration rate determined which regime was possible, with stronger migration rates favoring the latter. It is thus not surprising that we recover results similar to the ones that have been derived for when two of the three RI mechanisms interact. This implies that under the genetic architecture with polygenic adaptation and multiple DMIs considered here, it is unlikely that populations evolve towards build up of strong pre- and postzygotic barriers when there is gene flow. How loci involved in local adaptation and DMIs are distributed along the genome, and whether these different regimes lead to genetic signatures that can be detected based on polymorphism population genomics data remains to be further studied.

## Acknowledgments

We thank João Carvalho, Inês Fragata and the Evolutionary Genomics and Bioinformatics lab for discussion of the project and Claudia Bank for comments on the manuscript.

