## Appendix for "Interactions between mechanisms of reproductive isolation"

**Starting conditions:** We considered three possible starting conditions, corresponding to different biological scenarios: divergence from an ancestral (shared) state and secondary contact between already adapted populations that have accumulated genetic incompatibilities. The later case was investigated in the presence or absence of initial assortative mating. In details, we investigated:

- an "Ancestral" state, where both populations started identical and were fixed for allele 0 at all biallelic loci, resulting in the ancestral phenotype $Z_j = 0$, for each phenotypic trait $j$. There was no initial assortative mating ($P_C = 1$).
- a "Secondary contact" (or SC) state, where the two populations started with alleles differently fixed for each DMI locus, and at all remaining biallelic $L$ loci affecting the phenotype alleles took values such that individuals from each environment were close to their optimum (i.e., for each phenotypic trait $j$, $Z_j \approx Z_i^A$ for environment A and $Z_j \approx Z_i^B$ for environment B). There was no initial assortative mating ($P_C = 1$).
- a "Secondary contact with initial mate choice" (or SC + initial mate choice) state, where the starting conditions were identical to the "SC" state but with preexisting strict mate choice ($P_C = 10^{-4}$).

**Genetic architecture map:** A genetic architecture map was defined by the number of loci, the number of phenotypic traits they affected, their mutationnal effect on the phenotype, the number of DMIs, and the map of interacting loci. Unless specified otherwise, we used a single genetic background (i.e., map of genetic interaction and mutational effects) for a given set of relevant parameters (i.e., for $\{L, n_t, n_{\text{DMI}}\}$ ). In total, we investigated six different genetic architecture maps:

- "default" map: 50 DMIs pairs (100 loci, $n_{\text{DMI}} = 50$) among the $L = 500$ loci affecting a single phenotypic trait ($n_t = 1$). Each locus could be involved in at most a single DMI pair. This is the map illustrated in Figure 1B.
- "default 3D" map: 50 DMIs pairs (100 loci, $n_{\text{DMI}} = 50$) among the $L = 500$ loci affecting three phenotypic traits ($n_t = 3$). Each locus could be involved in at most a single DMI pair.
- "optimized" map: 250 DMIs pairs (500 loci, $n_{\text{DMI}} = 250$) among $L = 500$ loci affecting a single phenotypic trait ($n_t = 1$). The 250 loci affecting the phenotype positively interacted in a pairwise and unique way with the 250 loci affecting the phenotype negatively. We investigated this map, because, while unrealistic, it minimized the possible genetic conflict between ecological divergence and genetic incompatibilities, and therefore should be more likely to lead to accumulation of DMIs.

- "neutral" map: The genome was formed by $L = 800$ loci: half of them, chosen randomly, affected the phenotype, while the other 400 loci did not and formed 200 pairs of DMIs ($n_{\text{DMI}} = 200$). Each locus was involved in a single DMI pair. The phenotype consisted of a single trait ($n_t = 1$). We investigated this map to separate the loci involved in local adaptation and DMIs.

- "network" map: The genome was formed by $L = 800$ loci: half of them, chosen randomly, affected the phenotype, while 50 loci were randomly picked among the other 400 (neutral) loci and formed 200 DMIs pairs ($n_{\text{DMI}} = 200$). The phenotype consisted of a single trait ($n_t = 1$). We investigated this map to assess whether a network of DMIs promoted the accumulation of DMIs.

- "neutral display" map: 50 DMIs pairs (100 loci, $n_{\text{DMI}} = 50$) among $L = 500$ loci affecting the phenotype, which was defined as two phenotypic traits $n_t = 2$. In this case, the first phenotypic trait affected the extrinsic fitness and the second one was used as cue for mate choice. The mutational effects on the first phenotypic trait and the DMIs pairs were identical to the "default" map.

**Populations metrics during evolution:** We tracked the evolution of the different RI barriers per population by computing the mean phenotype for local adaptation, the mean choosiness for mate choice and the hybrid load between populations (defined below) for the accumulation of DMIs.

All metrics were computed, per population, for adult individuals after selection and migration had already occurred. In particular, this meant that if all individuals of the two populations were at the phenotypic optimum at A ($Z_j^A$) and B ($Z_j^B$), the mean phenotype for trait $j$ of population A depended on migration and was given by $\bar{Z}_j = (1 - m)Z_j^A + mZ_j^B$.

The hybrid load was defined as the reduction in (intrinsic) fitness caused by genetic incompatibilities. This was quantified by computing the intrinsic fitness component of simulated ephemeral (not part of the next generation) $\text{F}_1$ individuals resulting from the cross of individuals from the two populations, correcting by the mean intrinsic fitness to their parents ($\bar{w}_{int}^A$ and $\bar{w}_{int}^B$). This correction allows us to disentangle the hybrid load (defined as the reduction in fitness due to genetic incompatibilities present in different populations) from the mutational load caused by DMIs segregating within each population. The ephemeral $\text{F}_1$ individuals were obtained with males of population A (resp. B) choosing females of population B (resp. A), according to their choosiness. We focused on ephemeral $\text{F}_1$ individuals since all DMIs between populations would be in the double heterozygote state. Therefore, we could compute the number of equivalent fixed DMIs needed to generate the observed hybrid load (corrected by within population mutational load due to genetic incompatibilities):

$$n_{\text{DMI}}^{eq} = \frac{1}{log(1-\epsilon)} \times log\left(\frac{2\bar{w}_{F_1}^{int}}{\bar{w}_{int}^A + \bar{w}_{int}^B}\right) \tag{A1}$$

where $\bar{w}_{F_1}^{int}$ is the mean intrinsic fitness of hybrids, and $\epsilon$ is the strength of epistasis due to DMIs. In scenarios with migration, this equivalent number of fixed DMIs could be underestimated since populations would contain a proportion of individuals with the alternative phenotype. Hence, immigrant males, from population A, would prefer to mate with those females, with an environment A phenotype despite their lower extrinsic fitness. Such matings would produce individuals with the environment A phenotype, that were likely DMI-free, regardless of the genetic distance between the A and B population, reducing the generation of $F_1$s expressing DMIs, and therefore resulting in a lower hybrid load between populations. This effect could be further exacerbated by the dispersal stage, especially with "adult dispersal", when selection preceded migration.

**Measuring Reproductive Isolation (RI) post evolution** Defining and measuring RI between populations remains a debated topic (Westram et al., 2022). Here we followed the definition proposed by Westram et al. (2022) and measured RI as the reduction in gene flow at a neutral marker. Thus, we considered the fate of alleles at two neutral markers, one freely recombining and the other situated in the middle of the genome. One individual (referred to as focal) from the population A was transferred to population B. We followed the fate of the alleles from population A at these two neutral markers, which were introduced into population B with an absolute number of two allele copies at each locus. Measurement of RI were done after the last generation of the long term evolution. For convenience, we refer to this as post-evolution simulations, and its initial starting time as generation 0 post-evolution. The fate of these alleles was tracked with the two populations in isolation, regardless of the initial migration rate. Although invasion probability was the most obvious metric to track, it was computationally heavy, especially when considering large populations. Therefore, we measured the sojourn time (i.e., the time of loss of the neutral markers) or their frequency after $N$ generations post-evolution, if they were still polymorphic. We considered the fate of 1,000 invasion events (each time choosing the focal individual randomly) in populations of $N = 10,000$ individuals.

We measured the strength of RI post-evolution, accounting for the parameter values that evolved at each barrier (i.e., alleles at phenotypic loci, genetic incompatibility loci and male choosiness locus). To assess the relative impact of each mechanism of RI, we simulated the invasion of focal individuals either accounting for the effect of the three barriers, a combination of two barriers, or just one barrier. This was done by turning off barriers (regardless of whether the barrier barely or strongly evolved), which corresponded to fixing the extrinsic fitness, intrinsic fitness or male choosiness (or any combination of these two) to 1. For instance, to assess the relative impact of local adaptation, we computed the mean sojourn time by only considering the extrinsic fitness values ($w_{ext}^A$ and $w_{ext}^B$ at the two environments, respectively), setting the intrinsic fitness and male choosiness to 1 (i.e., $w_{int} = 1, P_C = 1$ ). Thus, there were seven different possible combinations of mechanisms of RI (referred to as "RI barriers" for the rest of manuscript). In details, the barrier could be formed by local adaptation alone ("LA"), mate choice alone ("MC") or the presence of DMIs alone ("DMIs"). It could also result from the effect of two mechansims ("LA+DMI", "LA+MC" or "MC+DMI") or of all three mechanisms ("all").

### Gene flow mostly caused by immigrant individuals of the choosy sex

Based on the mate choice scheme investigated, we expected that a male ("choice-making" individual) would have a higher chance to leave offspring after migration than a female ("non-choosing") individual, all things being equal. Indeed, a male, once selected as a potential mate based on its intrinsic and extrinsic fitness, only needed to accept one among $\chi$ potential mates, while a female, once chosen as a potential mate, had only one chance to be accepted for mating. Assuming all individuals were at the optima in each environment, this difference could easily

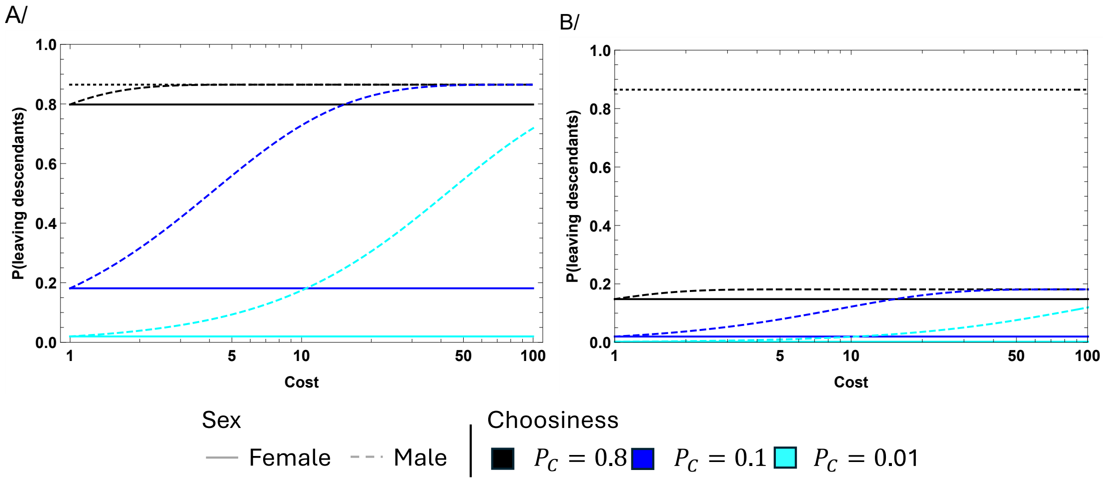


Figure A1: Probability of a focal immigrant individual (fitness $w_f$) to leave at least one descendant in a resident monomorphic population (fitness $w_r$) in the next generation ($P(desc.)$ as a function of the cost of choosiness $\chi$, in the absence (panel A, $w_f = w_r$) and presence (B, $w_f = 0.1$, $w_r = 1.0$) of local adaptation. We represent this probability for a focal male (dashed line; eq. A2) and female (eq. A3 solid line) and difference choosiness strength ($P_C = 0.8$ in black, $P_C = 0.1$ in blue and $P_C = 0.01$ in cyan). The probability for the neutral case is given by the dotted dashed line on both panels. Other parameters are $N = 1000$ and $w_r = 1$.

be quantified, considering that the resident population have a size $2N$ with 50:50 sex-ratio, all resident individuals have fitness $w_r$, and that the focal immigrant individual has fitness $w_f$. Under these assumptions, the probability that this focal individual left offspring in the next generation is, for a focal male:

$$P_m(desc.) = 1 - \left( \frac{(N-1)w_r}{w_f(1-(1-P_C)^{\chi}) + (N-1)w_r} \right)^{2N} \xrightarrow[N \to \infty]{} 1 - e^{2((1-P_C)^{\chi}-1)\frac{w_f}{w_r}} \tag{A2}$$

and for a focal female:

$$P_f(desc.) = 1 - \left( \frac{(N-1)w_r}{P_C w_f + (N-1)w_r} \right)^{2N} \xrightarrow[N \to \infty]{} 1 - e^{-2P_C\frac{w_f}{w_r}} \tag{A3}$$

where $P_C$ is the probability of a male accepting a mating with a female with an alternative phenotype, and $\chi$ is the number of potential mates considered by a male to reproduce with (details of the calculations are given in the SI section M6). These predictions matched simulations (Fig. S26), including under complete neutrality ($w_r = w_f = 1$, $P_C = 1$) and random mating.

Given these equations, immigrating focal females were far less likely to leave immediate descendants than their male counterpart, as the probability for females did not depend on the number of mates considered $\chi$ (Fig. A1). In contrast, the probability of leaving descendants for males increases as the number of mates considered $\chi$ increases. Thus, focal males and females are expected to have similar probabilities of leaving descendants only when the cost of choosiness is extremely high ($\chi = 1$). In contrast, under strong choosiness ($P_C \to 0$) and weak choosiness cost ($\chi \to \infty$), gene flow is expected to be sex-specific, mostly driven by the choosy sex. Since we fixed $\chi = 100$, in all our simulations gene flow was mediated mostly by males.
