## Supplement for "Interactions between mechanisms of reproductive isolation"

#### M1 Reformulation of Fisher's Geometric Model:

The general formulation of Fisher's geometric Model (Fisher, 1930), as presented by Gros et al. (2009), is given by:

$$w^A_{ext} = e^{-A_g\left(\Sigma_{j=1}^{n_t}(Z_j - Z_j^A)^2\right)^{\frac{Q}{2}}} \tag{S1}$$

with $n_t$ the number of phenotypic traits, $Z_j$ is the phenotypic value of trait $j$, $Z_j^A$ is the optimal phenotypic value for trait $j$ in environment A and $A_g$ a parameter scaling the average effect of mutations on fitness. However, such parameter do not have a direct connection to the model investigated here, and do not convey, in an intuitive way, how different the two environments are from each other. We noted that, when considering an individual perfectly adapted to environment B (i.e., $\forall j\ Z_j = Z_j^B$ ), we can write:

$$\underbrace{w^A_{ext}(B\ indiv)}_{\Omega^B} = e^{-A_g\left(\Sigma_{j=1}^{n_t}(\underbrace{Z_j}_{Z_j^B} - Z_j^A)^2\right)^{\frac{Q}{2}}} \tag{S2}$$

The sum term is simply the (square) distance in the phenotypic space between the optimum of the two environments, $\Delta_{AB} = \Sigma_{j=1}^{n_t}(Z_j^B - Z_j^A)^2$. Therefore, $\Omega^B$ can be an alternative parameter, to account for the difference between the A and B environment in our two-island model.

Therefore, the $A_g$ parameter can be written as function of the fitness of a perfectly adapted individual to one environment in the alternative environment $A_g = \frac{-log(\Omega_B)}{\Delta_{AB}^{Q/2}}$, assuming $\Delta_{AB} > 0$, leading to equation (1).

#### M2 F1 hybrid fitness

Based on equation (1), we could determine when $F_1$ individuals, from parents at the phenotypic optimum at the two environments (e.g., mother with phenotype $Z_j^A$ and father $Z_j^B$), displayed hybrid vigor (their fitness was greater than the mean of the parents' fitness; given by $\frac{1+\Omega^A}{2}$). Assuming that $F_1$ individuals had an intermediate phenotype (i.e., $\forall j, Z_j = \frac{Z_j^A + Z_j^B}{2}$, leading to $w_{ext}^{A,F_1} = e^{0.25^{Q/2} log(\Omega^B)} = (\Omega^B)^{2-Q}$; by symmetry $w_{ext}^{B,F_1} = (\Omega^A)^{2-Q}$), whether hybrid vigor was observed mainly depended on the mean epistasis $Q$ of the landscape (and the parents' fitness):

$$w_{ext}^{A,F_1} > \frac{1+\Omega^B}{2} \text{ if } Q > -\frac{\log\left(\frac{\log\left(\frac{\Omega^B+1}{2}\right)}{\log(\Omega^B)}\right)}{\log(2)} \tag{S3}$$

### M3 Reformulation of the choosiness function:

Similar to the expression of Fisher's Geometric model, mate choice can be written as a function of the phenotype of the male ($Z_j^m$) and female ($Z_j^f$) individuals:

$$P(mating) = e^{-A_m \Sigma_{j=1}^{n_t}(Z_j^m - Z_j^f)^2} \tag{S4}$$

with $A_m$ corresponding to a parameter scaling the average effect of mutations on fitness. As above, we note that we can write the probability of a successful mating between two individuals perfectly adapted to different environment, for let say a male perfectly adapted to environment A (with a phenotype $Z_j^m = Z_j^{A,m}$) and a female perfectly adapted to environment B (with a phenotype $Z_j^f = Z_j^{B,f}$):

$$P_C = e^{-A_m \underbrace{\Sigma_{j=1}^{n_t}(Z_j^{A,m} - Z_j^{B,f})^2}_{\Delta_{AB}}} \tag{S5}$$

Therefore the $A_m$ parameter can be written as $A_m = \frac{-log(P_C)}{\Delta_{AB}}$, assuming $\Delta_{AB} > 0$, leading to equation (3).

### M4 Mutational bias

Given that the range of values that the phenotype and mate choice could take was bounded, these traits suffered from a mutational bias: when the phenotypic trait (or the mate choice allele effect) reached values close to their extrema, the mean displacement caused by new mutations was, on average, no longer 0 but biased towards the other extremum (Fig. S27). Thus, mutational bias acts against divergence of populations and evolution of strict mate choice. This differs from other theoretical studies where mutation bias, in a male display trait, promoted the evolution of choosiness in females (Pomiankowski et al., 1991; Andersson and Iwasa, 1996). Furthermore, we considered assortative mating, whereas those theoretical studies considered a trait-preference model.

### M5 Probability of observing DMI between isolated population under the weak mutation strong selection approximation

To better understand how many DMIs we should expect between isolated populations, we considered the limiting case of weak mutation strong selection. For simplicity, we considered a haploid model, with 2 biallelic loci A and B. The haplotype *ab*, *Ab* and *aB* were viable (with fitness of 1) while haplotype AB was non-viable (fitness of 0). In addition, we assumed mutation happened at rate $\mu$ from ancestral to derive state, and at rate $\nu$ from derived to ancestral state. Under the weak mutation assumption, the fate of the new mutation would be resolved before a second one appeared. This corresponded to a Markov chain with the following transition rate

$$\begin{array}{cccc} & ab & Ab & aB \\ ab & 1-2\mu & \mu & \mu \\ Ab & \nu & 1-\nu & 0 \\ aB & \nu & 0 & 1-\nu \end{array} \tag{S6}$$

The stationary state of said chain was given by $P(Ab) = P(aB) = \frac{\mu}{2\mu+\nu}$ and $P(ab) = \frac{\nu}{2\mu+\nu}$, with $P(Ab)$ the probability that the population was fixed for haplotype $Ab$ at a given time. Given that the haplotype $AB$ is non-viable, we obtained $P(Ab) = P(A)$ and $P(aB) = P(B)$.

For symmetric mutation rate, it simplified to $P(A) = P(B) = \frac{1}{3}$. The probability of having a DMI between 2 populations ($p_1$ and $p_2$) was therefore given by $P(\text{DMI}) = \frac{2}{9}$. Indeed, $P(\text{DMI}) = P(\text{A in } p_1)P(\text{B in } p_2) + P(\text{A in } p_2)P(\text{B in } p_1) = 2 * P(\text{A in } p_1)^2$ or $2\left(\frac{\mu}{2*\mu+\nu}\right)^2$, since $P(\text{A in } p_1) = P(\text{A in } p_2) = P(\text{B in } p_1) = P(\text{B in } p_2)$.

#### M6 Probability of leaving descendants:

We consider a population of size $2N$, with $N$ females and $N$ males. For one of the sex, we have $N-1$ composed of individuals of type R and a single focal individual, of type F. Individuals are chosen to reproduce according to their fitness. If individuals are of the same type, they always produce an offspring. If they are of different types, they only produce an offspring with probability $P_C$. Males are chosen first, and can try to mate with up to $\chi$ females or until an offspring is produced.

First, we focus on the focal individual being a male. We define 3 random variables: $F$ tracking the resident vs immigrant status of the father (R for resident and I for immigrant), $X$ tracking whether the father was the first male considered (1 if it was the first or 0 if not), $Y$ tracking the resident status of the first considered male (R for resident and I for immigrant). Then, using the law of total probability and Bayes theorem we can derive the probability that a given offspring has a resident father $P_m(F = R)$:

$$\begin{aligned}
P_m(F=R) &= \Sigma_{x,y} P(F=R, X=x, Y=y) \\
P_m(F=R) &= P(F=R, X=1, Y=R) + \underbrace{P(F=R, X=0, Y=R)}_{=0} \\
&\quad + \underbrace{P(F=R, X=1, Y=I)}_{=0} + P(F=R, X=0, Y=I) \\
P_m(F=R) &= \underbrace{P(F=R, X=1 | Y=R)}_{=1} P(F=R) + P_m(R) \qquad \underbrace{P((X=0, Y=I) | F=R)}_{P(X=0,Y=I) \text{ since we randomly pick a new male}} \\
P_m(F=R) &= \frac{P(F=R)}{1-P(X=0,Y=I)} \\
P_m(F=R) &= \frac{P(F=R)}{1-P(Y=I)P(X=0|Y=I)}
\end{aligned} \tag{S7}$$

Each of these term can be easily computed with $P(F=R) = \frac{(N-1)w_R}{w_F+(N-1)w_R}$ (probability that the male chosen is a resident according to the fitness) , $P(F=I) = \frac{w_F}{w_F+(N-1)w_R}$ (probability that the male chosen is an immigrant according to the fitness) and $P(X=0|Y=I) = (1-P_C)^\chi$ (probability that the immigrant male reject all $\chi$ females).

As a consequence, we obtained:

$$P_m(F = R) = \frac{w_R(N-1)}{w_F(1-(1-P_C)^{\chi}) + w_R(N-1)} \tag{S8}$$

For females, we can notice that it is equivalent to the male case with the strongest cost of choosiness $\chi = 1$, and therefore the formula above applies. The probability that as least on offspring has the focal (or immigrant) male as its parent is given by $P_m(desc) = 1 - P_m(R)^{2N}$, leading to the equation (A2) and (when setting $\chi$ to 1) (A3) for females. Finally, it is important to remark that if choosiness differs between the 2 populations, for males it is the choosiness of the focal individual that matters, while for females, it is the choosiness of the resident males that do.

#### R1 Shape of ecological fitness landscape has reduced impact in the evolution of reproductive isolation barriers

We investigated whether the number of dimensions (i.e., number of traits) of the phenotypic space affected the evolution of the different mechanisms of reproductive isolation. As the number of phenotypic traits increased, the distance between the optimal phenotypes increased, therefore we set the position of the optima for each phenotypic trait such that the distance between optima at the two populations was constant, irrespective of the number of dimensions (for $n_t = 3$, the optimum for environment A is $(2.887, 2.887, 2.887)$ and $(-2.887, -2.887, -2.887)$ for B). Interestingly, we did not observe any significant changes in the evolution of reproductive isolation for all three mechanisms at different migration rates (Fig. S23).

We also investigated the shape of the fitness landscape. In principle, this parameter could have a large impact on the evolution of reproductive isolation, as it will impact the fitness of phenotypes in the vicinity of the optimum. When mean epistasis between mutations was positive ($Q < 2$), the fitness landscape became sharper, with small deviations in phenotype around the optimum leading to strong reductions in fitness. Oppositely, when mean epistasis was negative ($Q > 2$), the fitness landscape around the optimum became flatter (i.e., an individual with a phenotype away from the optimum can still have a high fitness). In addition, since the fitness of $F_1$ hybrid individuals between parents at the optima of the two environment (and therefore having an intermediate phenotype) is given by $w_{alt}^{2-Q}$, increasing $Q$ weakens the role of local adaptation in reproductive isolation. As illustrated in Figure S24 (for $m = 0.05$), the mean phenotype of the population sat further away from the optimum as $Q$ increased. Yet, strict mate choice evolved regardless of the mean epistasis in the fitness landscape. Finally, accumulation of DMIs was only observed for $Q < 2$ and only in some replicates.

To assess if our conclusions were affected by the inherent mutational bias in our model towards the phenotype value of zero for all traits (i.e., the same as the phenotypic value of $F_1$ hybrids for cases with symmetric optima), we repeated the simulations under models with asymmetric optima at the two populations. In the presence of local adaptation, we did not detect any changes due to the asymmetry (Fig. S25). The deviation of the mean phenotype from the optimum was the same regardless of the relative position of the ancestral phenotype and the environmental optima, indicating that the effect of selection and migration overshadow the effect of mutational bias.

#### R2 Evolution of RI in the absence of local adaptation

We investigated the consequence of the absence of local adaptation. Interestingly, we found that mate choice and DMIs could not maintain population differentiation in secondary contact models (Fig. S10). At high migration rate, the lack of differentiation was not caused by a collapse of RI, but through initial ecological exclusion of one of the two populations (Fig. S10, S21). Once one of the populations was lost, the role of mate choice and genetic incompatibilities became irrelevant and individuals in the two environments behaved as a single population. In contrast, at low migration rates ($m \leq 5 \times 10^{-4}$), we found that phenotypic differentiation converged to zero at similar rates for cases with and without migration, indicating that this is due to mutational bias (see Appendix). However, ignoring cases where ecological exclusion occurred, DMIs were actively purged with migration, as the hybrid load converged to zero (Fig. S10).

#### R3 Evolution of RI when mate choice is decoupled from the other barriers

For comparison with our results assuming a shared genetic basis for the three RI mechanisms, we considered an instance where the assortative mating was not decided upon a phenotype under local adaptation, but based on a "neutral display" phenotypic trait not affected by ecological selection. As we wanted to only consider the effect of one hypothesis, we maintain the pleiotropic nature of mutations, indicating that each mutation affected all phenotypic traits (whether or nor they were affect by ecological selection). Yet, mutational effects were independent between phenotypic traits, resulting in the independence between the two phenotypic traits. We therefore investigated the "neutral display" map with $n_t = 2$ phenotypic traits, where the first trait determined local adaptation (but not affecting mate choice) and the second trait used for assortative mating (but not under ecological selection). With assortative mating relying on a "neutral display" trait, mate choice was unable to evolve from an ancestral state nor from a secondary scenario (Fig. S11) although local adaptation evolved. Even when starting from secondary contact with initial mate choice (initial $P_C = 10^{-4}$), choosiness was slowly eroded, and populations converged to the same phenotypic value at the assortative mating trait. The loss of the DMIs always occurred first and seemed to be the trigger that led to the loss of mate choice. This was further confirmed by considering instances of increasingly costly DMIs (larger $\epsilon$): strict choosiness remained longer or persisted, as $F_1$s had a lower intrinsic fitness and hence were unlikely to be picked as potential mating partners (Fig. S22).

#### R4 Evolution of RI under different DMIs architectures

We further investigated whether the DMI architecture could explain the observed accumulation pattern. Indeed, this accumulation process can be strongly affected by whether incompatible loci are involved in a single or multiple DMIs (Guerrero et al., 2017). We thus compared the "neutral" architecture (loci forming DMIs were not affecting the phenotype) to the "network" architecture, where the same number of DMIs (200) was formed by only 50 loci. We found evolution towards lower hybrid load values for the "network" than the "neutral" architecture, suggesting that the lower the number of underlying DMI loci, the lower the number of equivalent fixed DMIs (Fig. S20B). In the "network" architecture, each locus was involved on average in 8 DMIs pair. If we consider a fixed derived allele, for the $i$ DMI pair, $A_i$ (in the ancestral background), then for the "neutral" architecture, its marginal fitness is reduced by the mutation from $b_i$ to $B_i$ at the incompatible locus. Therefore, mutation at the $B_i$ locus reduce the marginal fitness of allele $A_i$ by $2 * \mu(1-\epsilon)^2 + \mu^2(1-\epsilon)^4 \approx 2\mu\epsilon(2-\epsilon)$.Oppositely, in the "network" architecture, the $A_i$ loci interact on average with 8 $B_{i,j}$ loci. In the ancestral background, if allele $A_i$ is fixed, then mutation at $B_{i,1}$ locus will reduce the marginal fitness of allele $A_i$ by $\approx 2\mu\epsilon(2-\epsilon)$. Mutations at locus $B_{i,2}$ will also have the same effect (and up to locus $B_{i,8}$). Overall, marginal fitness of allele $A_i$ was reduced by a factor $\approx 8$, and therefore selection caused by background mutation and epistasis prevented the fixation of derived alleles, even in the absence of gene flow. Indeed, under the "network" architecture, we did not find an accumulation of DMIs, even in scenarios without migration.

We also investigated a DMI architecture map, which should favor the accumulation of DMIs. Indeed, as detailed in the Appendix, 250 loci affecting the phenotype positively interacted in a pairwise and unique way with the 250 loci affecting the phenotype negatively. For this "optimized" architecture, despite the fact that DMIs were fully correlated with the phenotypic effect of alleles, the proportion of equivalent fixed DMI evolved towards values similar to the "default" architecture ($\approx 0.15$). Importantly, with migration, accumulation of DMIs was strongly limited in all investigated DMI architectures, especially for more complex networks of interaction, suggesting that local adaptation did not facilitate the accumulation of DMIs.

### Supplementary figures

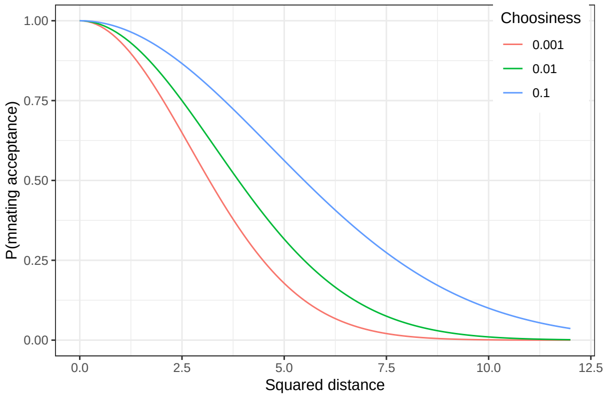


Figure S1: **Shape of the probability of accepting a mating depending on $P_C$ as a function of the distance between possible mates.**

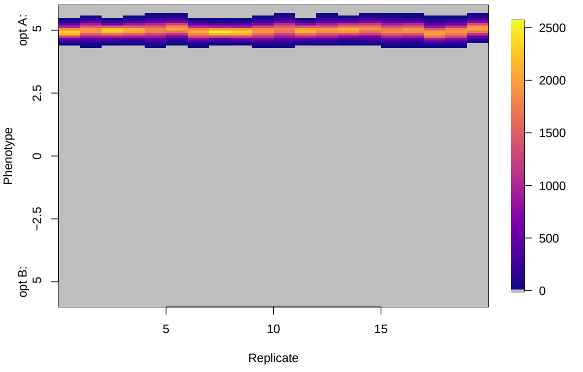


A/ Distribution of phenotype in isolated populations

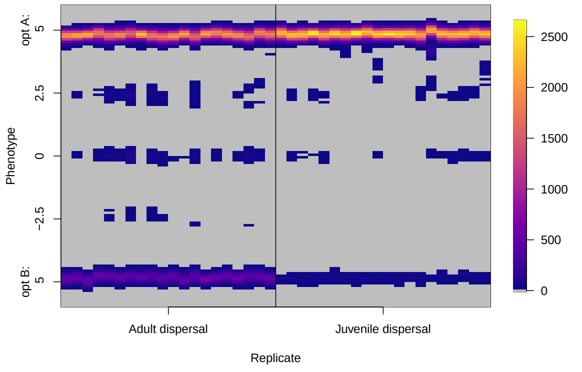


B/ Distribution of phenotypes in populations with migration ($m = 0.2$)

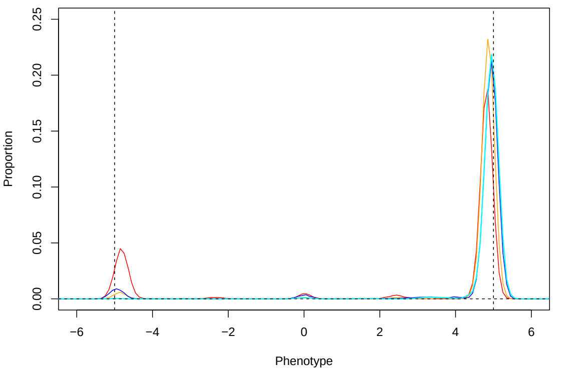


C/ Mean distribution of offspring phenotype, over 10 replicates and for different migration rates

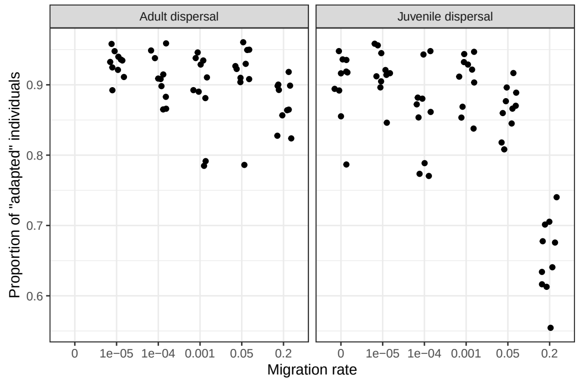


D/ Proportion of “fully” adapted individuals

Figure S2: **Local adaptation occurs across the migration rates considered.** Distribution of offspring phenotype (i.e. before migration) in isolated populations (A/ $m = 0$), or with strong migration (B/ $m = 0.2$) after 50N generation. For $m = 0.2$, we represent the distribution of offspring phenotypes for both life cycles “Adult dispersal” (left) and “Juvenile dispersal” (right) in the population residing in the A environment (optimal phenotype $z_O^A = 5$), for 20 replicates. The color scale corresponds to the absolute frequency. C/ Mean distribution of males offspring phenotype (i.e. before migration) with default (blue and cyan, $m = 0.05$) and strong (red ad orange, $m = 0.2$) migration rate after 50N generation across 10 replicates. We represent the mean distribution of offspring phenotypes for both dispersal stages, “adult dispersal” (blue and red) and “juvenile dispersal” (cyan and orange), in the population residing in the A environment (optimal phenotype $z_O^A = 5$). D/ Proportion of individuals that are “fully” adapted depending on the migration rate and the life cycle, defined as individuals whose phenotypes falls within the $5^{th}$ and $95^{th}$ percentile of the distribution of phenotype evolved in isolation computed over 10 replicates (i.e, between 4.64 and 5.32). The different panels correspond to the last time point of displayed in Figure 2 (more precisely, Figure 2 displays the mean phenotypes of the parents, while we focused here on the phenotype of their offspring - before selection or migration). Other parameters correspond to the default values.

A/ Isolated population ($m = 0$)

| | 0 | 0.1 | 0.2 | 0.3 | 0.4 | 0.5 | 0.6 | 0.7 | 0.8 | 0.9 | 1 |
|---|---|---|---|---|---|---|---|---|---|---|---|
| 0 | 35 | 246 | 47 | 25 | 17 | 19 | 34 | 18 | 30 | 50 | 385 |
| 0.005 | 0 | 315 | 40 | 26 | 8 | 20 | 10 | 4 | 14 | 8 | 126 |
| 0.010 | 0 | 21 | 0 | 0 | 0 | 0 | 0 | 0 | 0 | 0 | 0 |
| 0.015 | 0 | 2 | 0 | 0 | 0 | 0 | 0 | 0 | 0 | 0 | 0 |

B/ Weak migration ($m = 0.0005$)

| | 0 | 0.1 | 0.2 | 0.3 | 0.4 | 0.5 | 0.6 | 0.7 | 0.8 | 0.9 | 1 |
|---|---|---|---|---|---|---|---|---|---|---|---|
| 0 | 9 | 228 | 38 | 18 | 22 | 11 | 20 | 12 | 31 | 32 | 318 |
| 0.005 | 0 | 416 | 52 | 25 | 18 | 12 | 7 | 11 | 11 | 20 | 139 |
| 0.01 | 0 | 42 | 1 | 0 | 0 | 0 | 0 | 0 | 0 | 0 | 0 |
| 0.015 | 0 | 6 | 0 | 0 | 0 | 0 | 0 | 0 | 0 | 0 | 0 |
| 0.02 | 0 | 1 | 0 | 0 | 0 | 0 | 0 | 0 | 0 | 0 | 0 |

C/ Default migration ($m = 0.05$)

| | 0 | 0.1 | 0.2 | 0.3 | 0.4 | 0.5 | 0.6 | 0.7 | 0.8 | 0.9 | 1 |
|---|---|---|---|---|---|---|---|---|---|---|---|
| 0 | 7 | 203 | 44 | 18 | 26 | 10 | 24 | 23 | 25 | 45 | 268 |
| 0.01 | 0 | 386 | 53 | 27 | 13 | 18 | 18 | 11 | 16 | 26 | 188 |
| 0.02 | 0 | 21 | 0 | 0 | 0 | 0 | 0 | 0 | 0 | 0 | 3 |
| 0.03 | 0 | 9 | 1 | 0 | 0 | 0 | 0 | 0 | 0 | 0 | 1 |
| 0.04 | 0 | 3 | 0 | 1 | 0 | 0 | 0 | 0 | 0 | 0 | 0 |
| 0.05 | 0 | 2 | 1 | 0 | 0 | 0 | 0 | 0 | 0 | 0 | 0 |
| 0.06 | 0 | 4 | 2 | 0 | 1 | 0 | 1 | 0 | 0 | 1 | 0 |

Table S1: **DMIs were often polymorphic within population, but the rarer incompatible allele (almost) never exceed 5%.** Distribution of the derived allele per pair of DMIs, with the locus with the higher frequency of the derived allele given as column and the frequency of the derived allele at the rarer locus given as rows. Frequency were always round up to the next category to make sure that the 0 category truly indicates a lack of the derived allele. This was computed for population A at generation $50N$ and for different migration rates (A/ $m = 0$, B/ $m = 0.0005$ and C/ $m = 0.05$) and summed over 30 replicates. In total, 39.6%, 50.7% and 53.8% of all pairs of DMIs had derived mutations at the two incompatible loci for respectively $m = 0$, $m = 0.0005$ and $m = 0.05$.

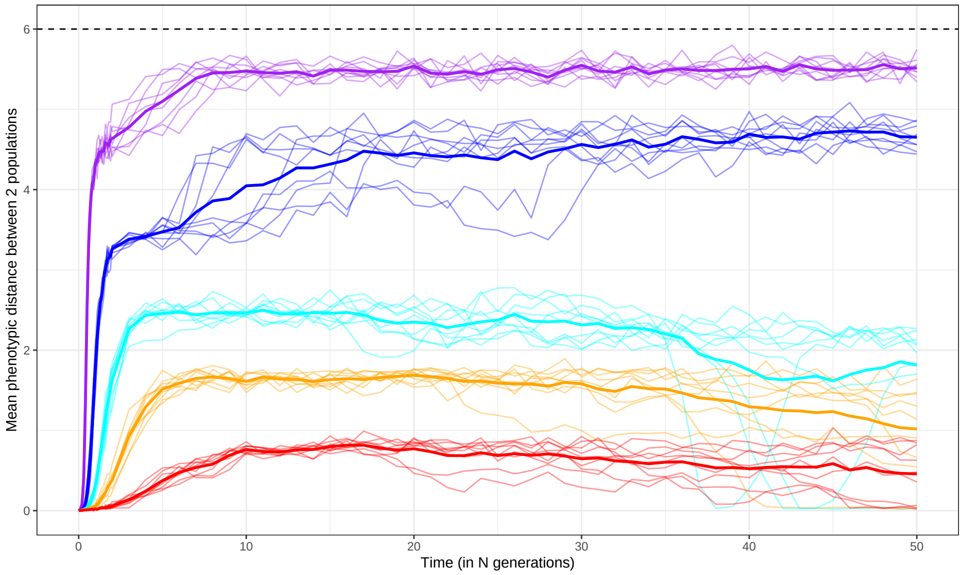


Figure S3: **Populations can diverge even when the fitness difference between F1 and optimum are less than the migration rate.** Evolution of the phenotypic distance between the two populations for different strength of selection ($w_{alt} = 0.1$ in purple, $w_{alt} = 0.2$ in blue, $w_{alt} = 0.3$ in cyan, $w_{alt} = 0.4$ in orange and $w_{alt} = 0.5$ in red) with strong migration ($m = 0.2$). Importantly, these values translate to the following extrinsic fitness for F1 individuals: $w_{F1} = 0.562$, $w_{F1} = 0.669$, $w_{F1} = 0.74$, $w_{F1} = 0.795$ and $w_{F1} = 0.841$. Life cycle is "juvenile dispersal". Other parameters correspond to the default values.

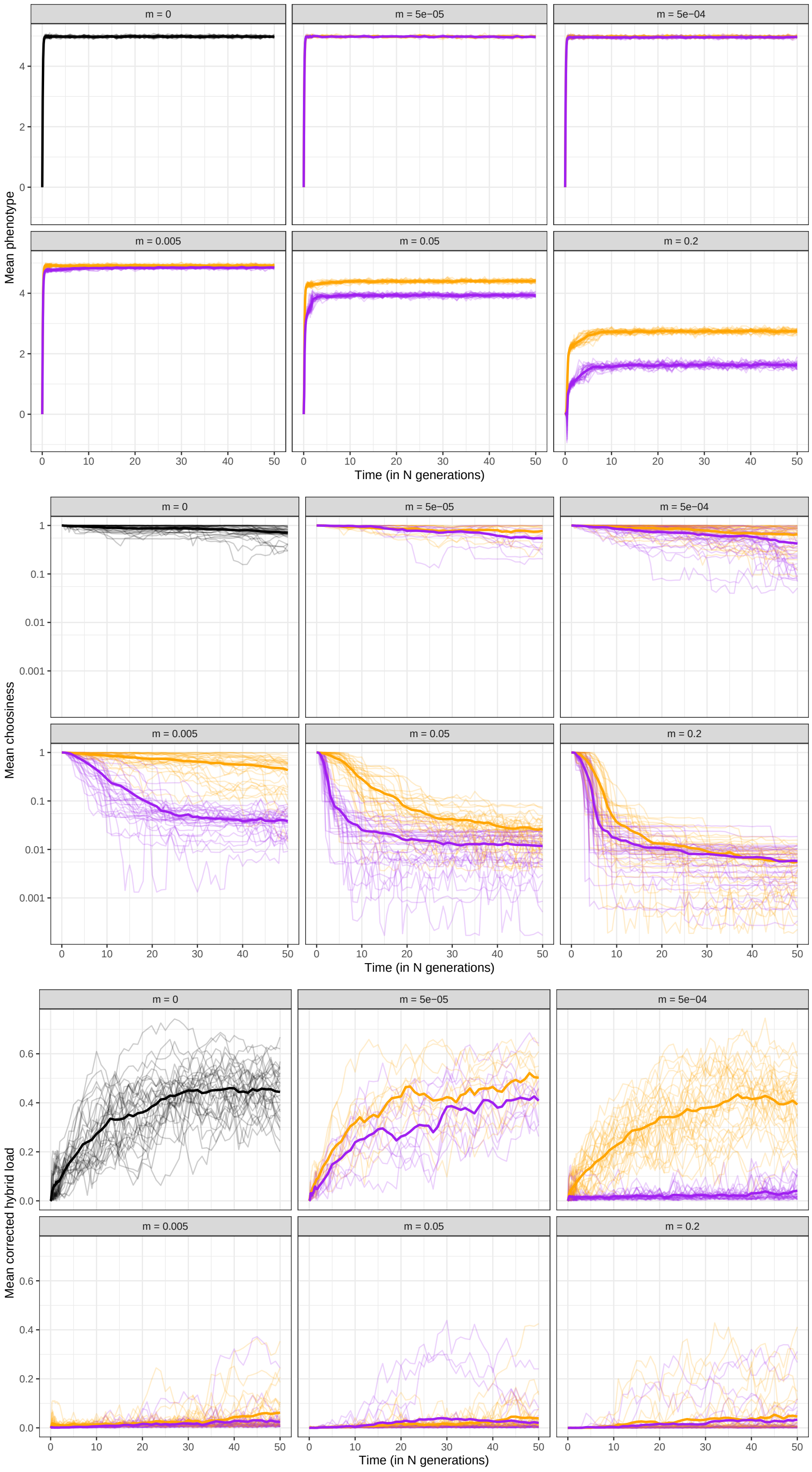


Figure S4: **Evolution under "default" genetic architecture of phenotypic traits (affecting local adaptation and mate choice) and genetic incompatibilities.** Evolution of the phenotype (top), mate choice (middle; on a log scale) and corrected intrinsic hybrid load (bottom) for different migrations rate (given in the facet header) and different life cycle ("juvenile dispersal" in orange and "adult dispersal" in purple). Each thin line corresponds to a different replicate, and the thick line to the mean over the 30 replicates (except for $m = 5.10^{-5}$, with only 10 replicates). The genetic map used is the "default" one and other parameters correspond to the default values and are given in Table 1.

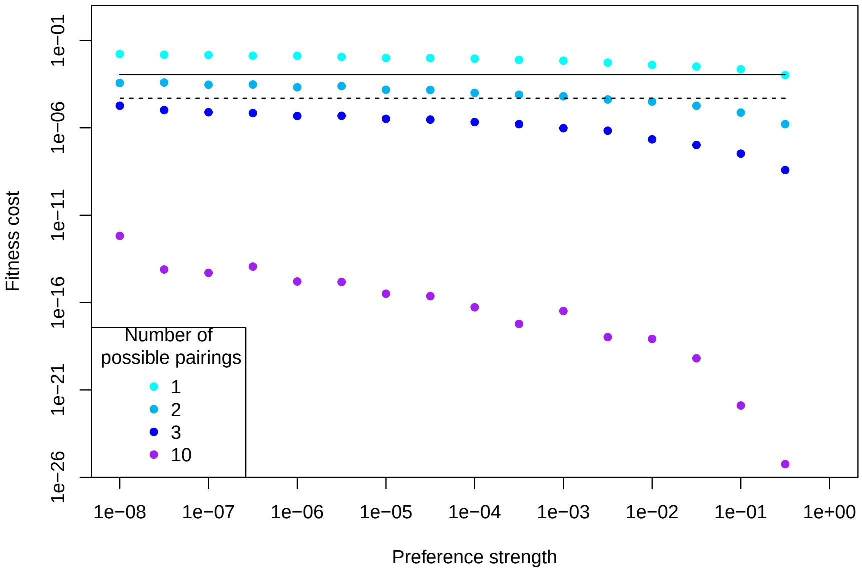


Figure S5: **The reduction in fitness caused by choosiness is weaker than the effect of drift except if the number of potential mates considered is really small ($\chi < 3$).** The color indicates different cost of choosiness (i.e, the number of mates considered by the choosing individual). The fitness reduction caused of choosiness is computed as the probability of an individual failing to find mate, given a choosiness and cost of choosiness values. For the choosy individual, it means rejecting $\chi$ potential mates, while for the non-choosy one, it means being rejected by one mate. Therefore, this cost ($s_c$) is given by $\prod_{k=1}^{\chi} 1 - P(mating) = s_c$, with $P_C$ given in equation (3). The $\chi$ possible mates were selected according to their fitness (sampling with replacement). The black dashed line corresponds to the effect of drift $\frac{1}{2N}$. The black line corresponds to the mean mutational load of the population, caused by new mutations, standing genetic variation and recombination, moving the next generation around the optimum phenotype. The distribution of phenotypes and the mutational load were directly taken from simulations done in the absence of mate choice, DMIs and migration. We used 1,000 simulations to compute the reduction in fitness caused by choosiness. Other parameters correspond to the default value and are given in Table 1

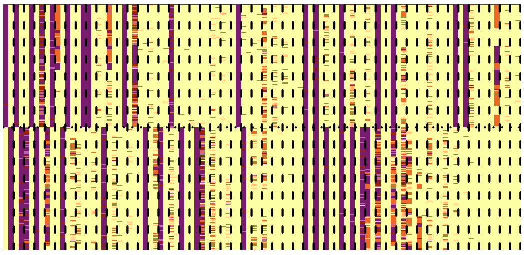

A/ $m = 0$

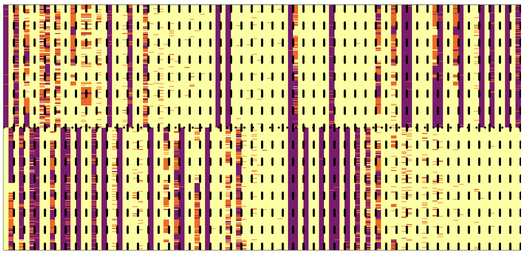

B/ $m = 5 * 10^{-5}$

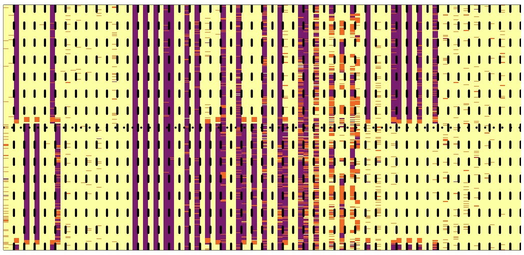

C/ $m = 0.05$

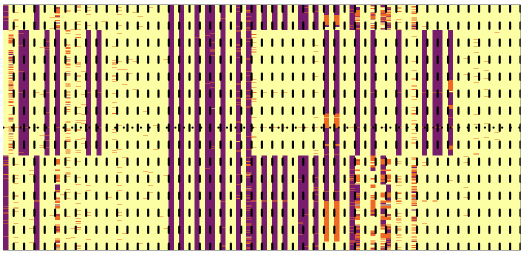

D/ $m = 0.2$

Figure S6: **DMIs were removed from the population when there was migration.** We displayed here the genetic maps of all 100 DMI loci, for different migration rates and for a subset of 500 individuals per population, for replicate 1. Each locus, invovled in a DMI, corresponded to a column, and each individual to a row. The colours indicated the genotypes at the DMI, with the homozygous ancestral state in yellow, the heterozygous state in orange and the homozygous derived state in purple. Interacting loci were displayed next to each other, with dashed lines separating each DMI pair. The top part corresponds to population 1 and the bottom one to population 2, separated by the horizontal dotted black line. A genetic incompatibility that could be expressed in F1 would correspond to a case where the two populations had different genotypes in each column of a given DMI (e.g. having purple on one diagonal and yellow on the other). As can be seen, as migration rates increased, for most DMI pairs the two populations fixed the same allele at each locus. The DMIs were ordered by the strength of the potential interaction: $f_1(A)f_2(B) + f_1(B)f_2(A)$ and the individuals are ordered via clustering using Manhattan distance. For migration $m = 0$ (A/), there were accumulation of DMIs, with the first 5 pairs of DMIs responsible for most of the post-zygotic isolation. Life cycle is “adult dispersal”. Other parameters correspond to the default values.

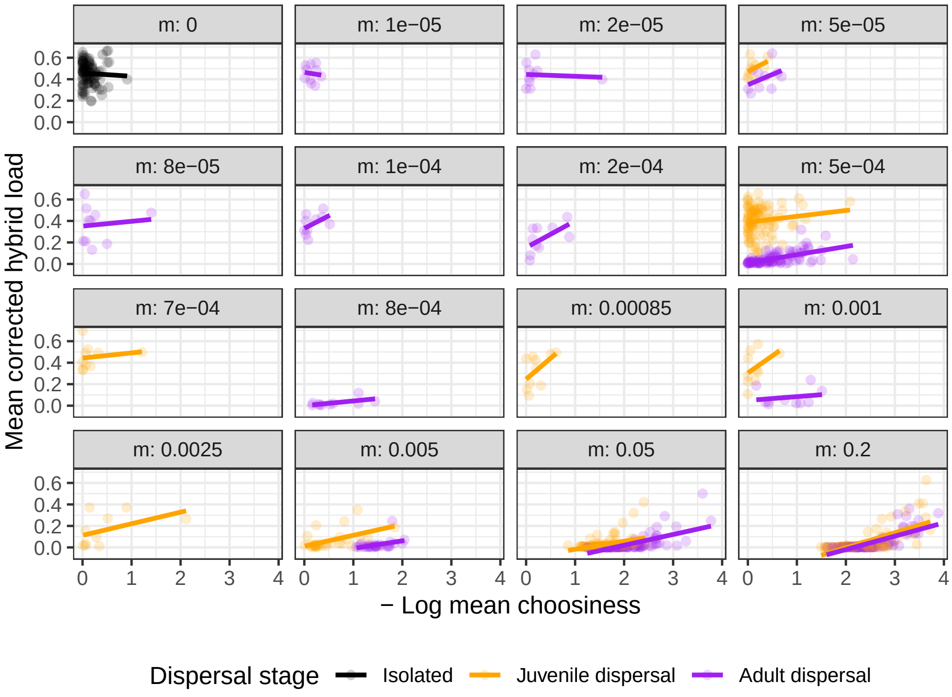


Figure S7: **Evolution of strict mate choice correlates with stronger hybrid load** We displayed here the correlation between (-log) mate choice (strict mate choice corresponds to large values) and hybrid load (right panel) at the end of the $50N$ generations for both life cycles ("Adult dispersal" in purple and "Juvenile dispersal" in orange) and different migration rates (given in the facet header). The lines corresponds to the linear regression between log mate choice and hybrid load for the replicates of a given parameter combination. Other parameters correspond to the default value and are given in Table 1.

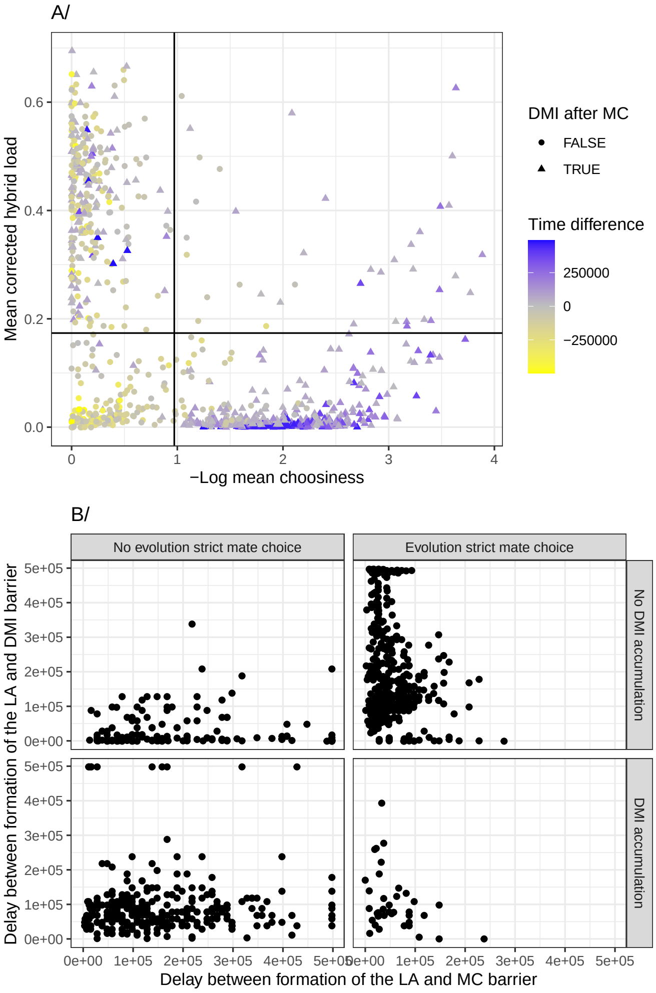


Figure S8: **Co-occurence of hybrid load and mate choice is due to DMIs accumulation after the evolution strict mate choice** A/ Relation between (-log) choosiness (strict mate choice corresponds to large values) and hybrid load at the end of the $50N$ generations. Color and shape indicates the delay between the formation of the mate choice and accumulation of DMIs barrier (we used both here to better visualize between values close to 0) B/ Comparison of the delay between the formation of the local adaptation and the mate choice barrier to the delay between the formation of the local adaptation and the accumulation of DMIs barriers. Formation of the barrier was defined as reaching 75% of the maximum mean value at generation $50N$. Facet corresponds to whether the barrier did actually evolved, as defined previously. The points displayed here matched the one in Figure 3. The genetic map used was the “default” one, other parameters correspond to default values given in Table 1.

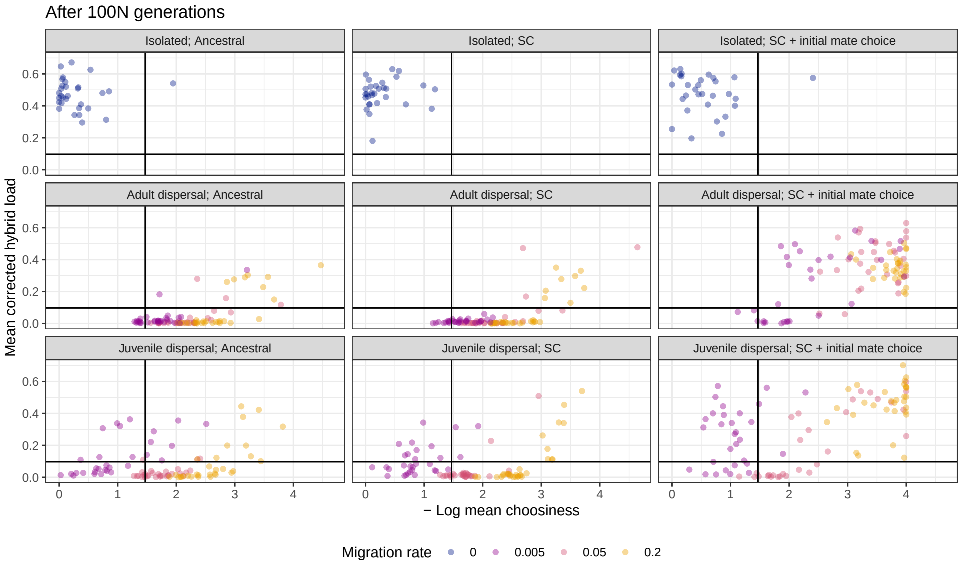


Figure S9: **Accumulation of DMIs with non-negligible migration generally required extremely strict mate choice ($P_C < 10^{-3}$)** Link between $F_1$ hybrid load and (-log) mate choice (strict mate choice corresponds to large values) for different starting conditions and migration rates. Each points corresponds to generation $100N$ of a different replicate (30) and corresponds to the final state of the evolutionary trajectories displayed in Figure 4. The black lines corresponded to the thresholds defined in the main text: the minimum of mean mate choice and hybrid load among the 100 isolated replicates starting from an ancestral population. Other parameters correspond to the default value and are given in Table 1

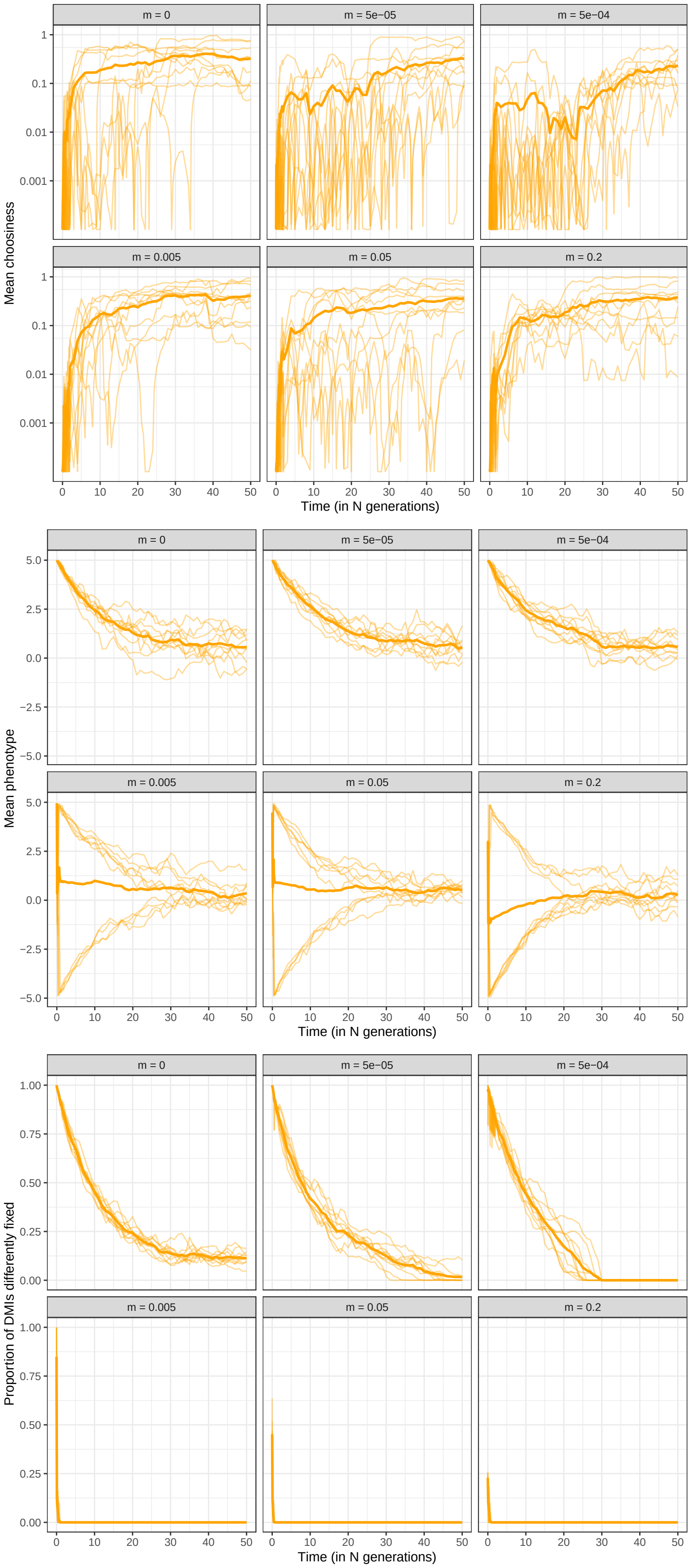


Figure S10: **Upon secondary contact and without local adaptation, and strong RI barriers, population differentiation is lost due to ecological exclusion.** Evolution of mate choice (top; on a log scale), phenotypic trait (middle) and F1 hybrid load (bottom) starting with divergence with incompatibilities and strict initial mate choice, in the absence of local adaptation and for different migration rate. Migration rates are given above each panel. Other parameters correspond to the default value and are given in Table 1

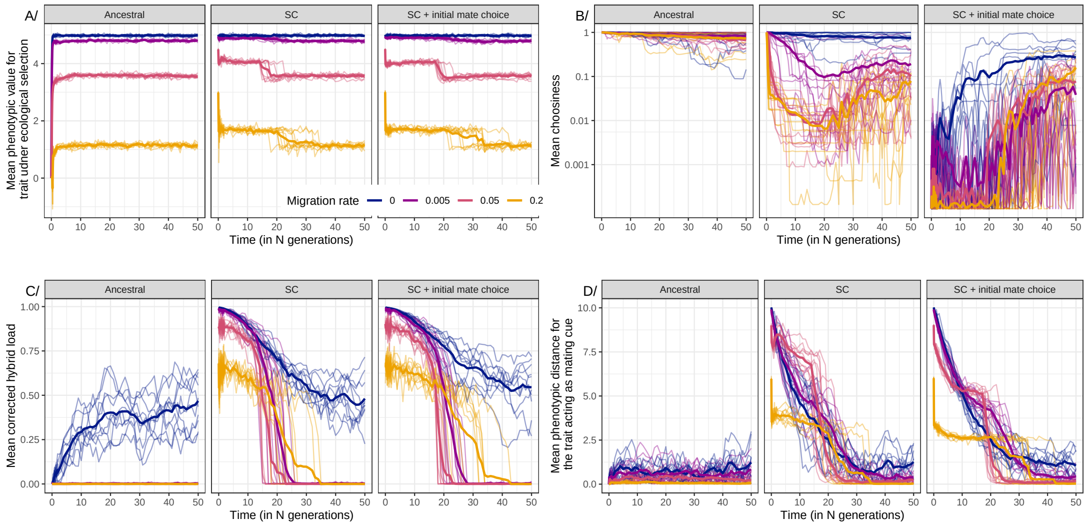


Figure S11: **Strict mate choice does not evolve is the "mating cue" is not a magic trait.** Evolution (in population A) of the first phenotypic trait (under ecological selection, panel A), mate choice (panel B; on a log scale), corrected hybrid load (panel C) and phenotypic distance between population at the trait acting as mating cue (not under ecological selection) and for different starting conditions (given in the facet header - panel D). Different colors correspond to different migration rate. The solid lines corresponds to the mean over the 10 replicates. The genetic map used was the "neutral display", other parameters correspond to default values given in Table 1.

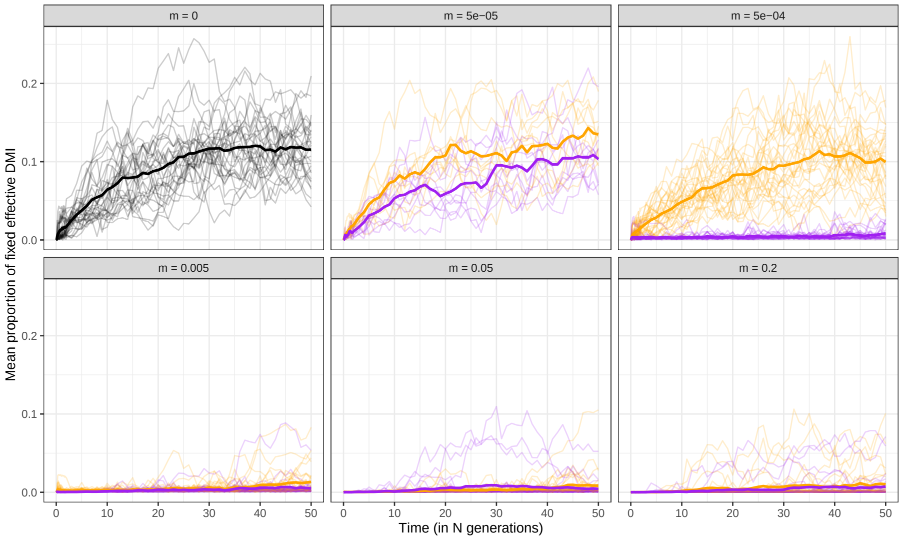


Figure S12: **Migration dictated whether DMIs accumulated, but the proportion of DMIs fixed between populations converged to similar values.** Evolution of the mean proportion of effective DMIs (as defined by equation (A1). For different migration rates (given in the facet header) and different life cycle ("juvenile dispersal" in orange and "adult dispersal" in purple). Each thin line corresponds to a different replicate, and the thick line to the mean over the 30 replicates. The genetic map used is the "default" one and other parameters correspond to the default values and are given in Table 1

.

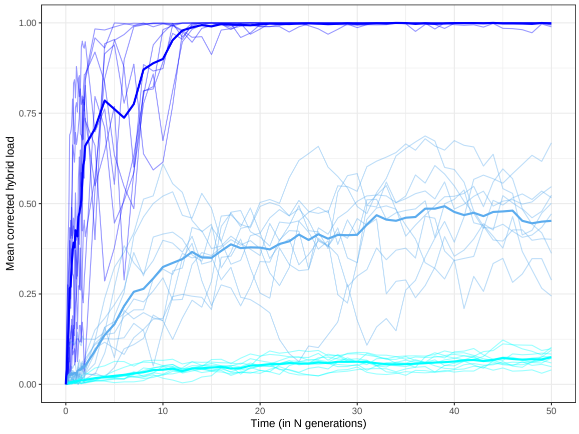


A/ Mean corrected hybrid load

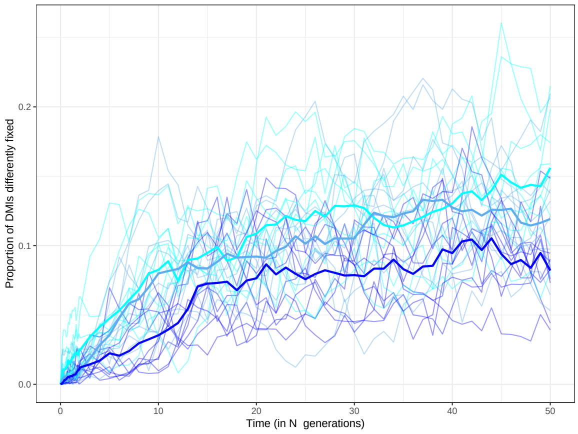


B/ Mean proportion of differently fixed effective DMIs

Figure S13: **Strength of epistasis had weak affect on the proportion of fixed effective DMIs between populations** Evolution of the intrinsic hybrid load (top) and the effective number of DMIs (bottom) for different strengths of the DMIs (cyan $\epsilon = 0.01$, blue $\epsilon = 0.1$, dark blue $\epsilon = 0.9$) in isolation. Each thin line corresponds to a different replicate, and the thick line to the mean over the 10 replicates. The genetic map corresponds to the "default" map and other parameters correspond to the default value and are given in Table 1

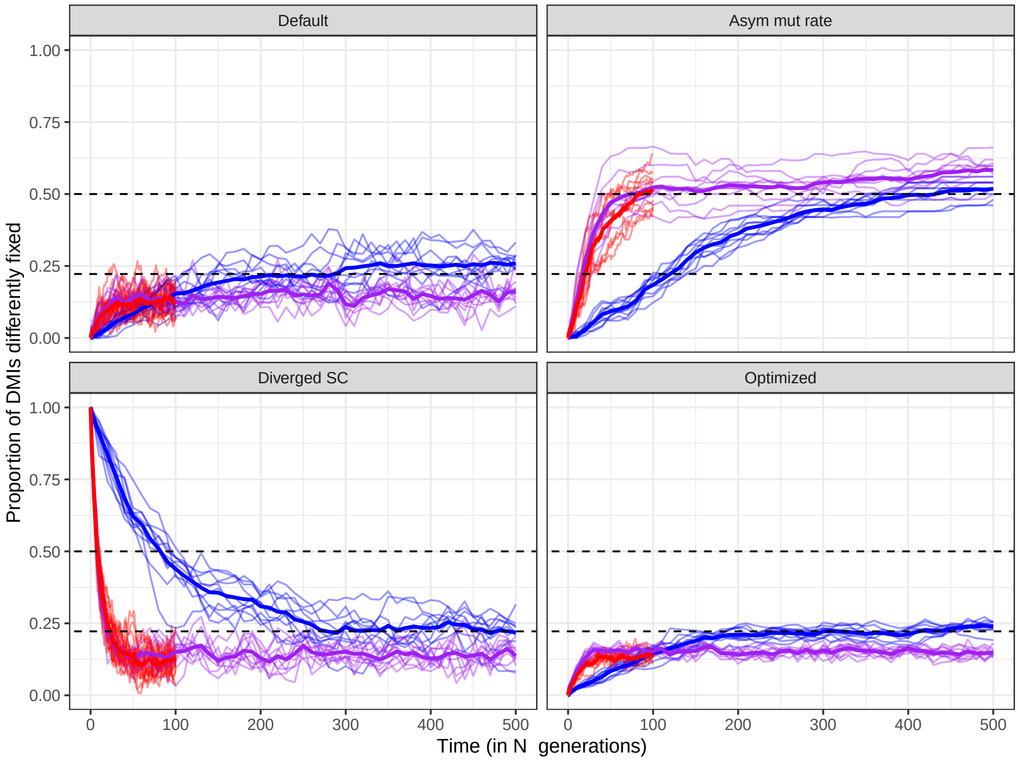


Figure S14: **Symmetry in mutation process and scaled mutation rate influenced the proportion of effective DMIs fixed between isolated populations.** Proportion of differentially fixed pairs of effective DMIs for isolated populations, for different population size and values of the mutation rate (($N = 1000, \theta = 0.001$) in blue, ($N = 1000, \theta = 0.01$) in purple and ($N = 10000, \theta = 0.01$) in red). The black dashed lines corresponded to the weak mutation strong selection approximation, derived in the main text, for symmetric (2/9) mutation rates or in the absence of back mutation (1/2). Each facet corresponded to a different starting condition and present one differences from the "Default" case. "Default" corresponded to populations diverging an ancestral one, a symmetric forward and backward mutation rate and using the "default" genetic map. For "Diverged SC", the populations started not in the ancestral state but being already locally adapted and having all 50 DMIs differently fixed. For "Asym mut rate", the backward mutation rate (from derived to ancestral) was 100 times lower than the forward rate (kept to $\mu = 10^{-6}$). For "Optimized", we used the "optimized" genetic map instead, where all loci formed pairwise DMIs, with always an allele increasing the phenotypic value interacting with one decreasing it. For each combination of parameters, the replicates are displayed in thin lines, while the average given by the thicker line. The number of fixed effective DMIs is computed from the mean hybrid load (see equation (A1)). Other parameters correspond to the default value and are given in Table 1.

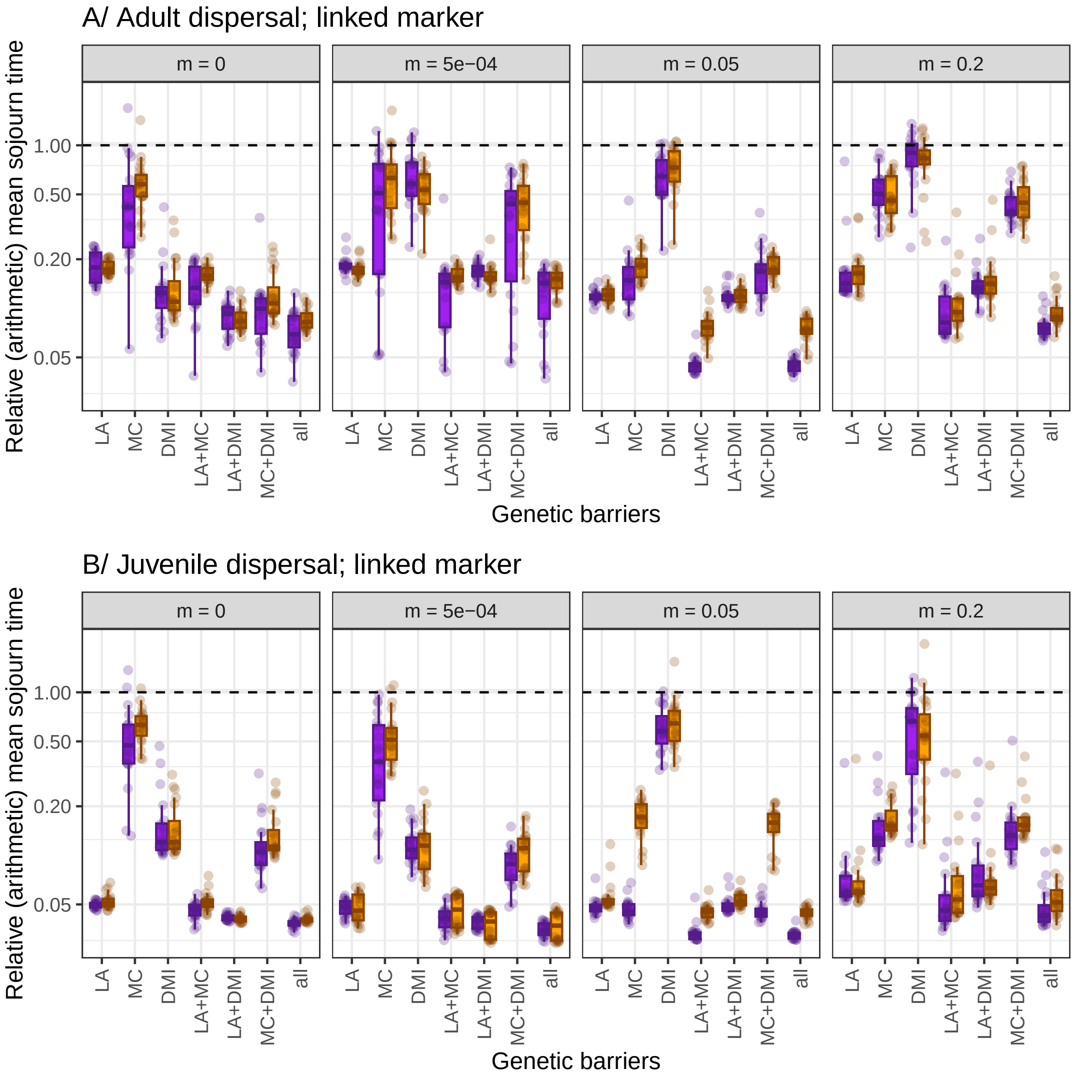


Figure S15: **Linked markers experience stronger reduction in sojourn time, but the patterns remain qualitatively similar to the ones observed for unlinked marker** Relative sojourn time of a linked marker introduced by an immigrating individual from the alternative population with different reproductive barriers for males (orange) and females (purple). The sojourn time is measured relative to either the neutral cases with the normalizing factor being the mean sojourn time over replicates. Panel A corresponds to the “adult dispersal” life cycle and panel B to the “juvenile dispersal” one. Other parameters correspond to the default values.

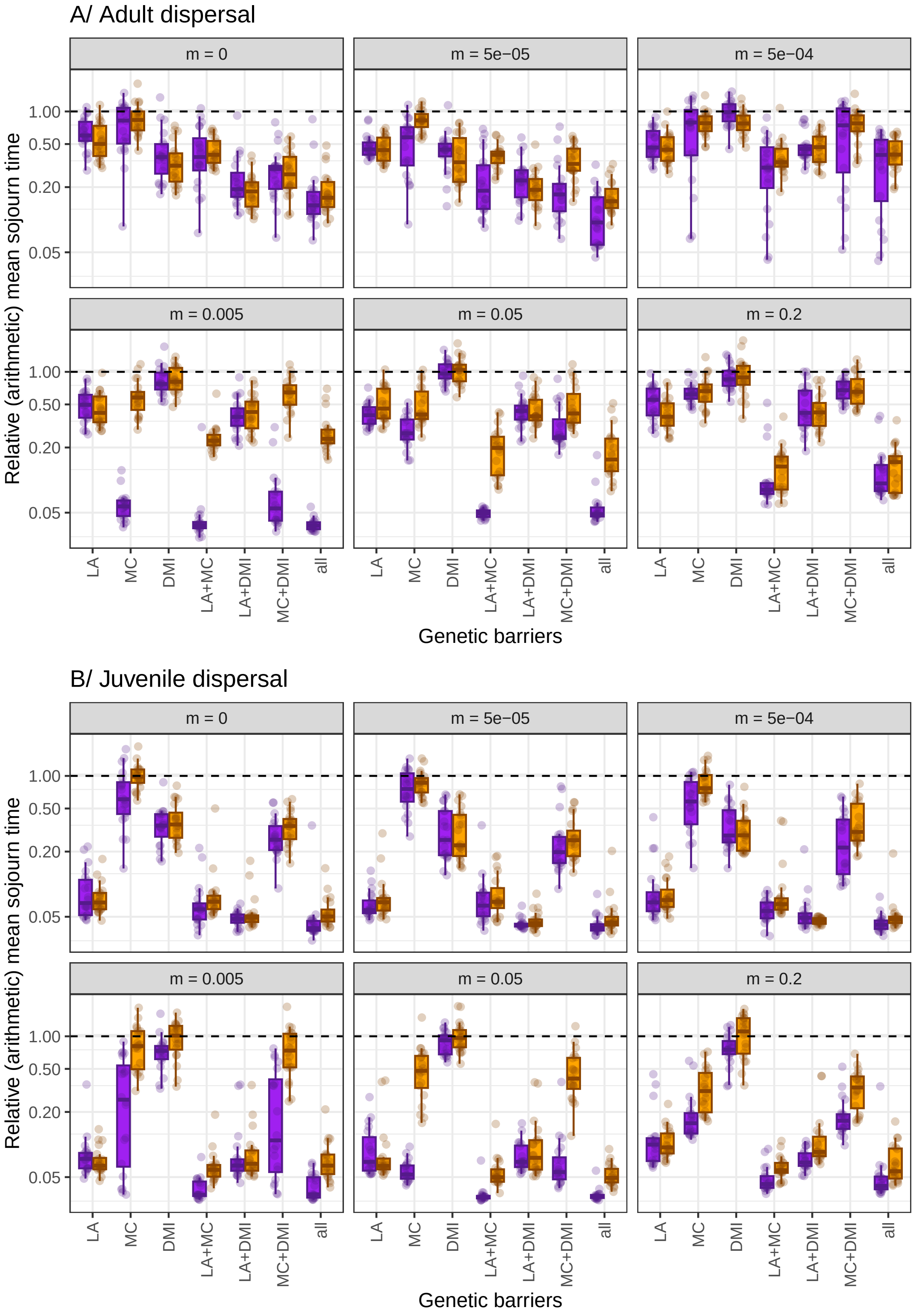


Figure S16: **Intermediate migration rate lead to the evolution of the strongest "all" barrier with "adult dispersal".** Relative sojourn time of a freely recombining marker introduced by an immigrating individual from the alternative population with different reproductive barriers for males (orange) and females (purple). The sojourn time is measured relative to either the neutral cases with the normalizing factor being the mean sojourn time over replicates. Panel A corresponds to the "adult dispersal" life cycle and panel B to the "juvenile dispersal" one. This figure is identical to Figure 6, but with 2 additional migration rates ($m = 5 \times 10^{-5}$ and $m = 5 \times 10^{-3}$). The underlying absolute values are given in Figure S17. Other parameters correspond to the default values.

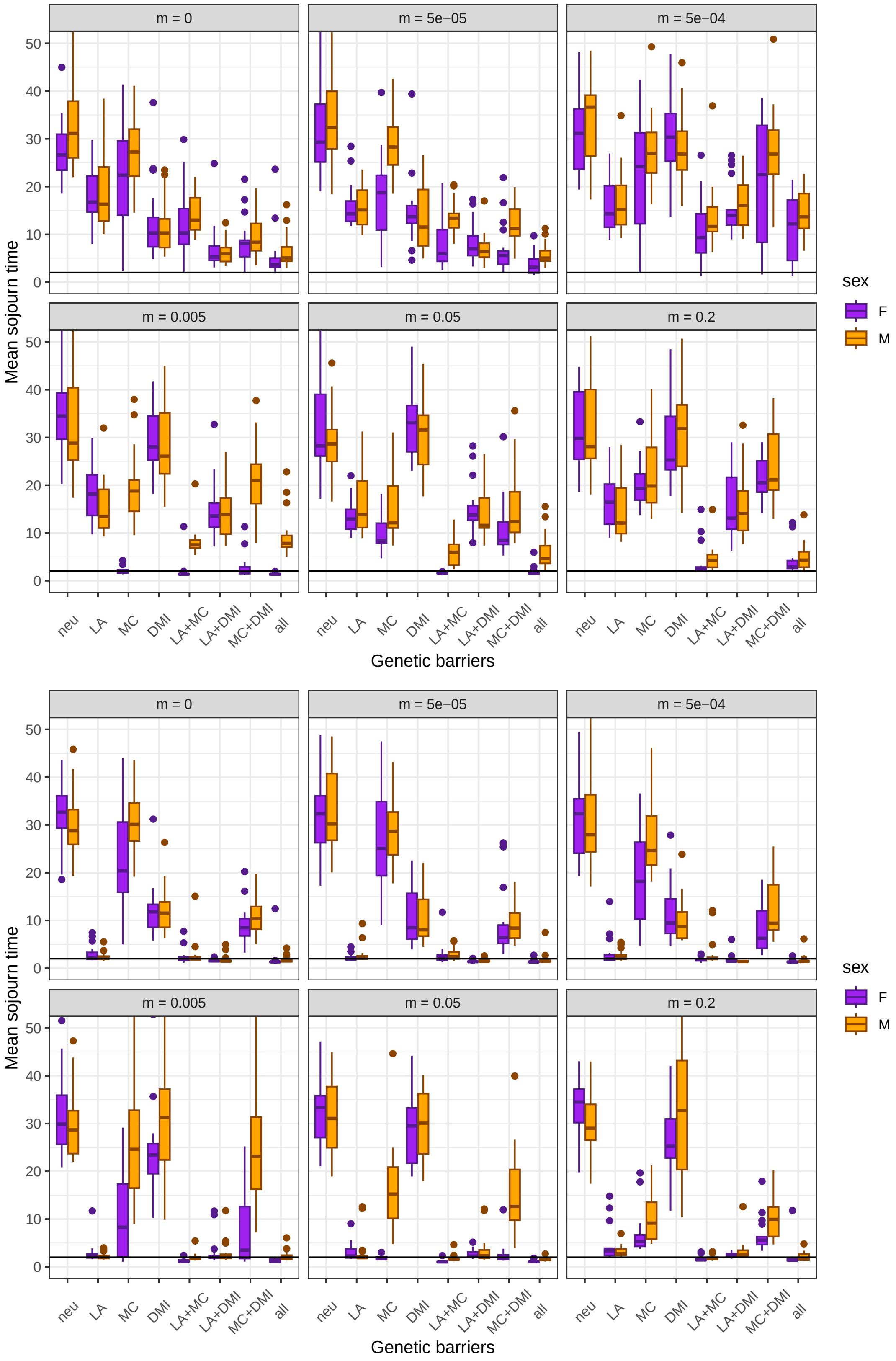


Figure S17: **Absolute sojourn time of a freely recombining marker** The sojourn time is measured by introducing by an immigrating individual from the alternative population with different reproductive barriers for males (orange) and females (purple). The top panel corresponds to the "adult dispersal" life cycle and the bottom panel the "juvenile dispersal" one. The black lines corresponds to a sojourn time of 2 generations (below this value immigrant focal individual died did not leave descendants). Points above 50 are not displayed for readability. Other parameters correspond to the default values.

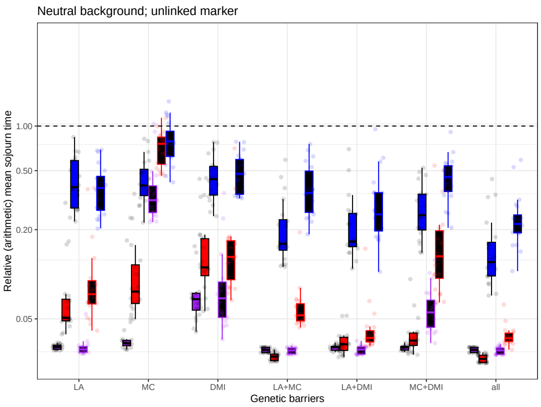


A/ "Juvenile dispersal"

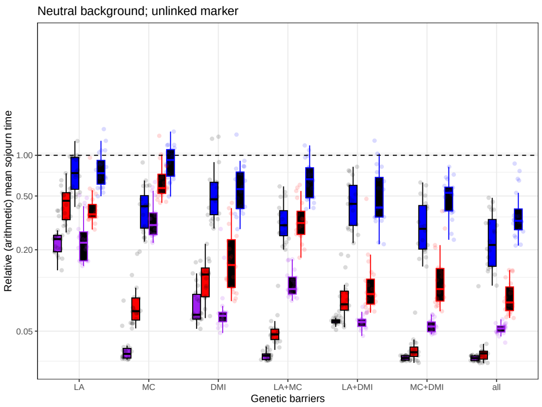


B/ "Adult dispersal"

Figure S18: **Local adaptation is a strong but brittle reproductive isolation barrier.** Relative mean sojourn time of an unlinked marker, when introduced through a female (colored background - black frame) or male (black background - colored frame) individual, with panel A corresponding to the "adult dispersal" life cycle and panel B to the "juvenile dispersal" one. Color indicates the strength of the barriers: blue 0.5, red 0.1, and purple 0.01. Life cycle is "juvenile-dispersal". Other parameters correspond to the default values.

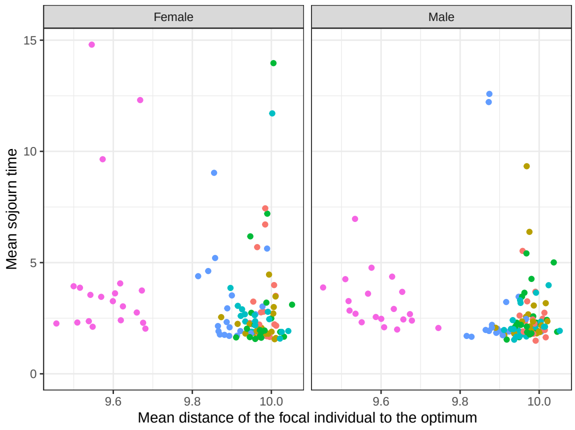


A/ Effect of local adaptation

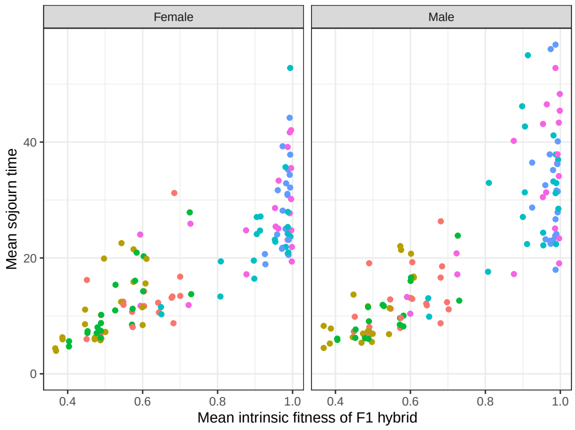


B/ Effect of the accumulation of DMIs

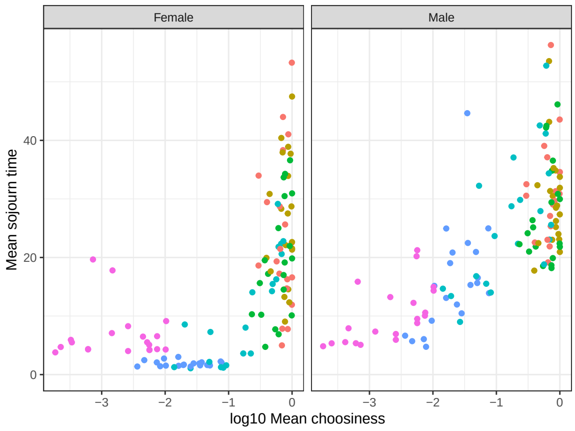


C/ Effect of mate choice

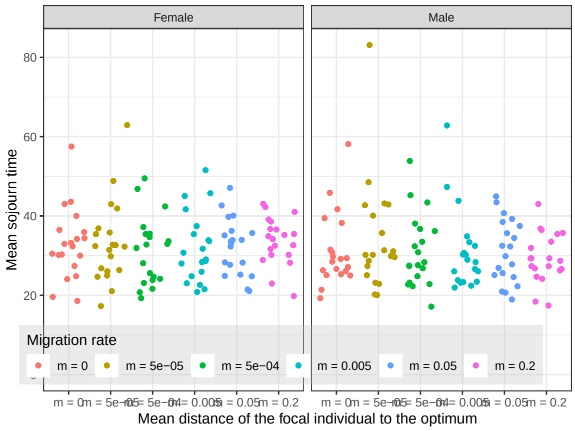


D/ No RI mechanisms

Figure S19: **Effect of the different RI barriers (A/ Local adaptation, B/ Accumulation of DMIs, C/ Mate choice or D/ in their absence) on the absolute sojourn time of an unlinked marker for the "juvenile dispersal" life cycle.** The x-axis corresponds for panel A/ to the mean distance between the focal individual and the phenotype optimal of the environment, for panel B/ to the mean intrinsic fitness of F1 hybrid, for panel C/ the mean choosiness and for panel D/ to the migration rate. The means were computed over the 1,000 invasion events. Color indicates the migration rates. Other parameters correspond to the default values.

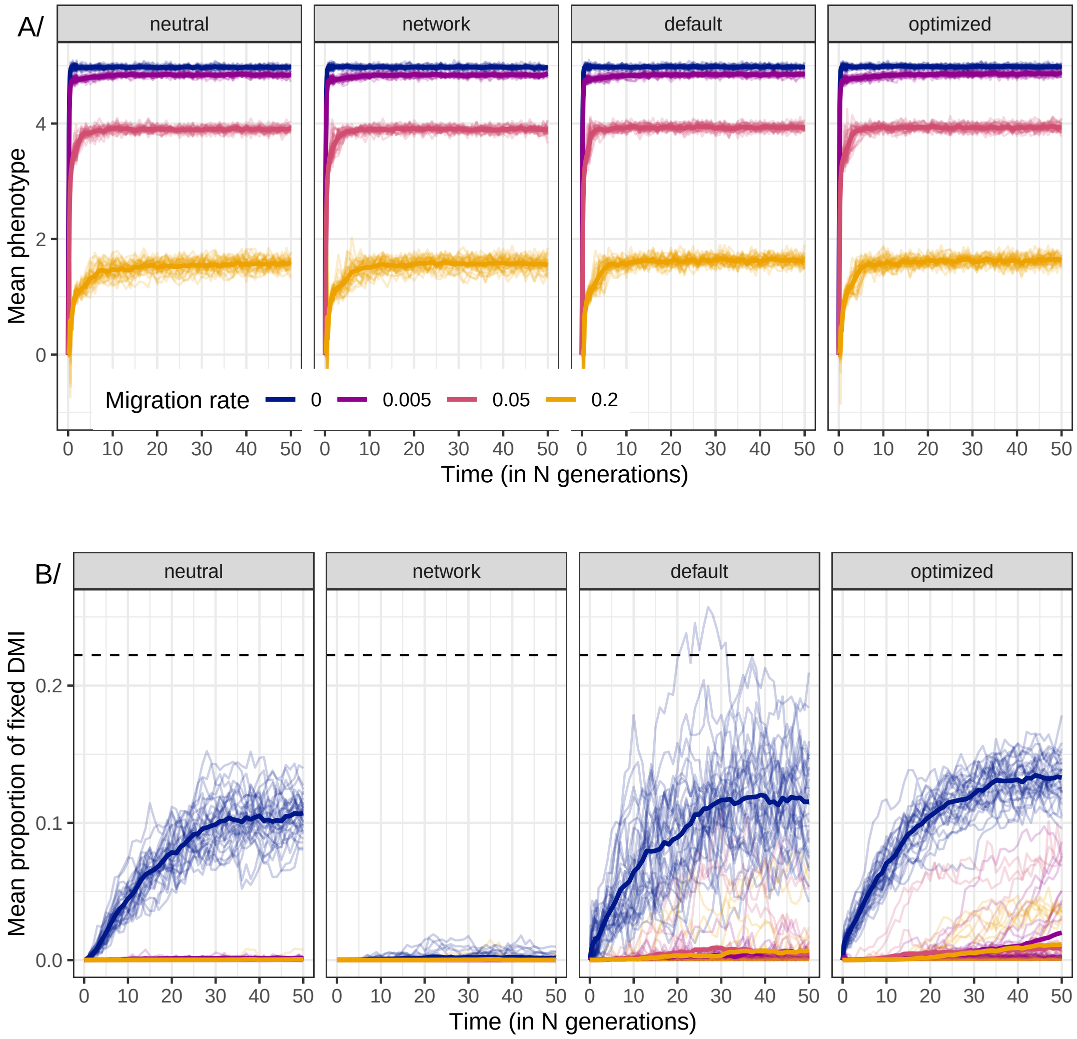


Figure S20: Evolution (in population A) of the phenotype (panel A) and number of equivalent fixed DMIs (B; computed according to equation (A1)) for different DMIs architecture (given in the header) and different migration rates. Each thin line corresponds to a different replicate, with the thick line corresponding to the mean over the 30 replicates. Color corresponds to different migration rates: $m = 0$ in blue, $m = 0.005$ in purple, $m = 0.05$ in pink and $m = 0.2$ in orange. For the B panel, the black dashed line corresponds to the weak mutation strong selection approximation for the proportion of DMI fixed between populations, $\frac{2}{9}$. The "neutral" architecture corresponds to 400 loci, not affecting the phenotype, forming 200 pairs of DMIs, "default" network corresponds to the 100 loci out of 500 forming 50 pairs of DMIs, the "network" one to 50 loci out of 800 forming 200 pairs of DMIs, and the "optimized" one to 500 loci out of 500 forming 250 pairs of DMIs, with always an allele increasing the phenotypic value interacting with one decreasing it (see methods). Other parameters correspond to the default values and are given in Table 1

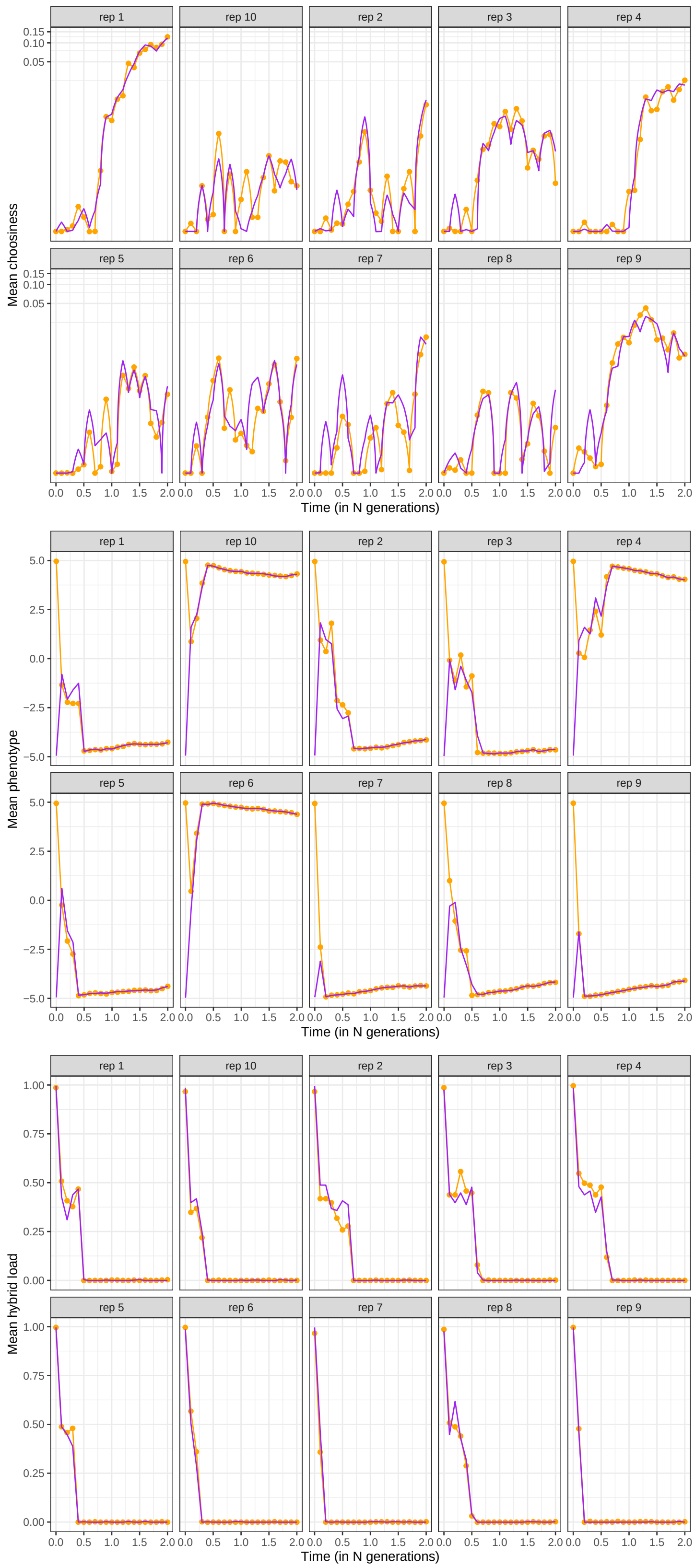


Figure S21: **Ecological dynamics responsible for the lack of differentiation between population.** Evolution of mate choice (top; on a log scale), phenotypic trait (middle) and F1 hybrid load (bottom) starting with divergence ($Z_1 \approx 5$ for individuals in population 1 and $Z_1 \approx -5$ in population 2), with incompatibilities and strict initial mate choice, in the absence of local adaptation ($w_{ext} = 1$). This figure is a detailed version of Figure S10 for $m = 0.005$ and only for the first $2N$ generations, with each panel corresponding to a different replicate. Population 1 is displayed in orange (line and points for visibility) and population 2 in purple. Epistasis strength is given by $\epsilon = 0.105$. Other parameters correspond to the default value and are given in Table 1

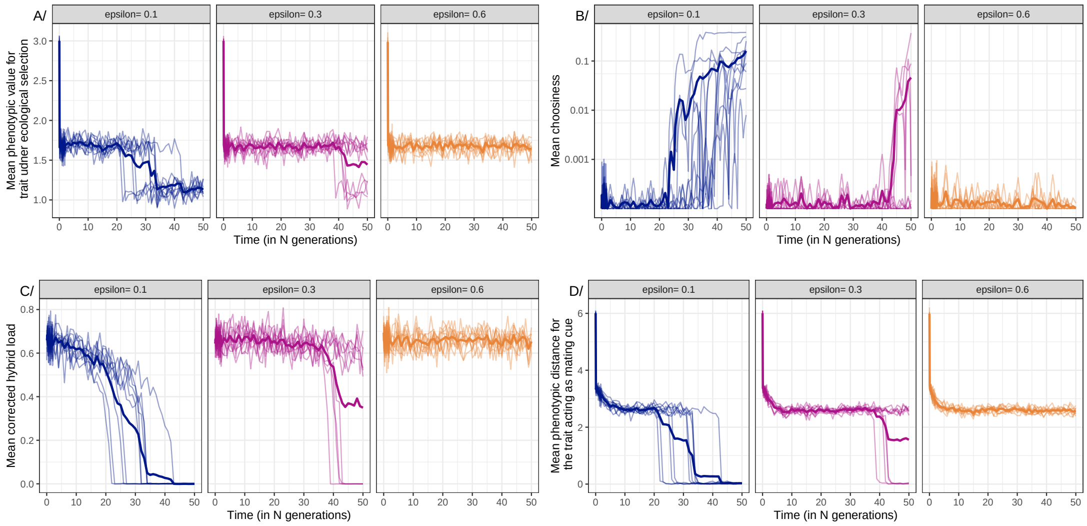


Figure S22: **Strong post-zygotic allowed the persistence of mate choice when the "mating cue" is not a magic trait** Evolution (in population A) of the first phenotypic trait (under ecological selection, panel A), mate choice (B; on a log scale), hybrid load (C) and (D) phenotypic distance for the assortative mating trait (not under ecological selection) for different strength of intrinsic incompatibilities ($\epsilon$, given in the facet header) and with strong migration ($m = 0.2$). The solid lines corresponds to the mean over the 10 replicates. The genetic map used was the "neutral display", and the starting conditions "Secondary contact with initial mate choice"; other parameters correspond to default values given in Table 1.

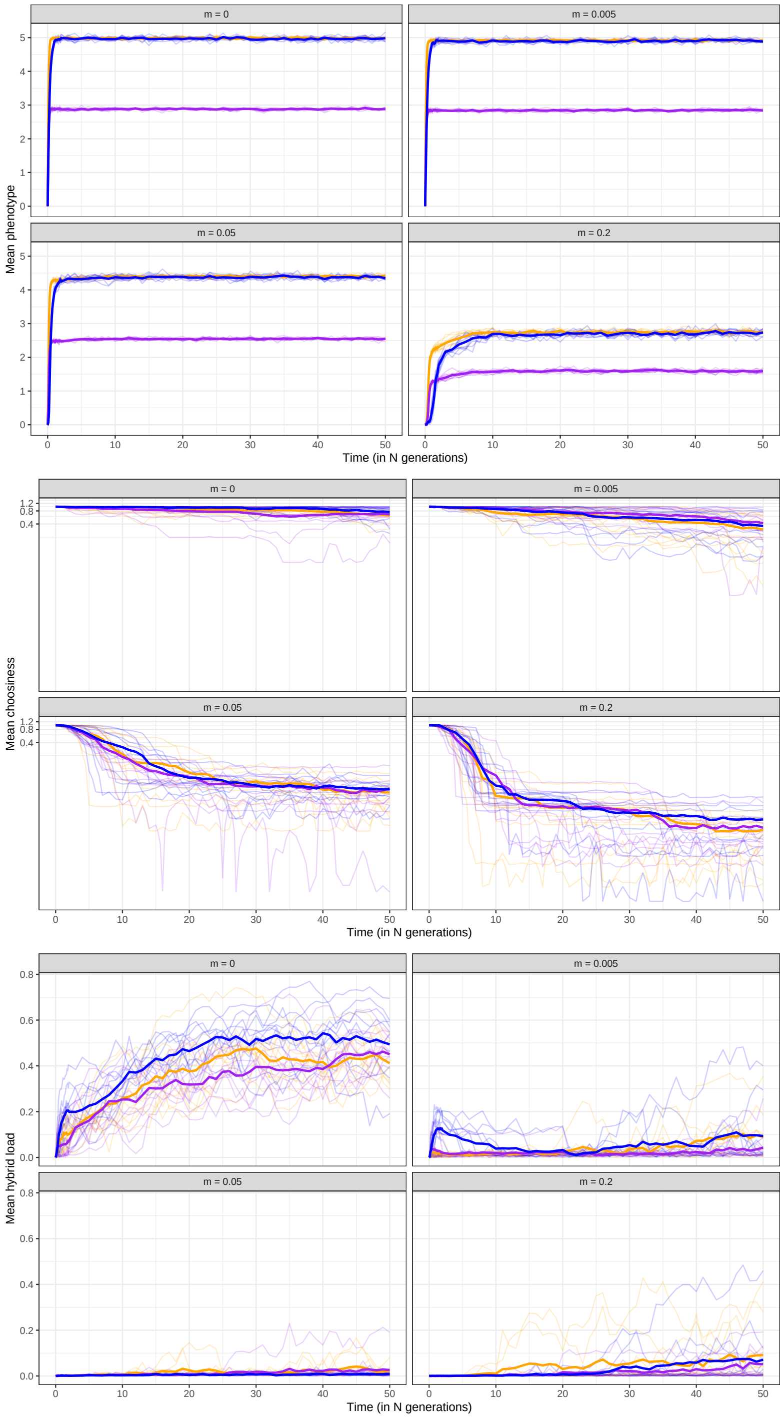


Figure S23: **Number of phenotypic dimension has limited impact on the evolutionary dynamics** Evolution of the phenotype (top), mate choice (middle; on a log scale) and intrinsic hybrid load (bottom) for different migrations rate (given in the facet header) and different number of phenotypic dimensions: orange for $n_{dim} = 1$ and $\Delta_{AB} = 10$, red for $n_{dim} = 3$ and $\Delta_{AB} = 17$, purple for $n_{dim} = 3$ and $\Delta_{AB} = 10$, blue for $n_{dim} = 3$ and $\Delta_{AB} = 10$ but with an asymmetric position of the optimum ($Z_j^A = 6.774$ and $Z_j^B = 1$). Each thin line corresponds to a different replicate, and the thick line to the mean over the 10 replicates. The genetic map corresponds to the "default 3D" map, life cycle is "juvenile dispersal" and other parameters correspond to the default value and are given in Table 1

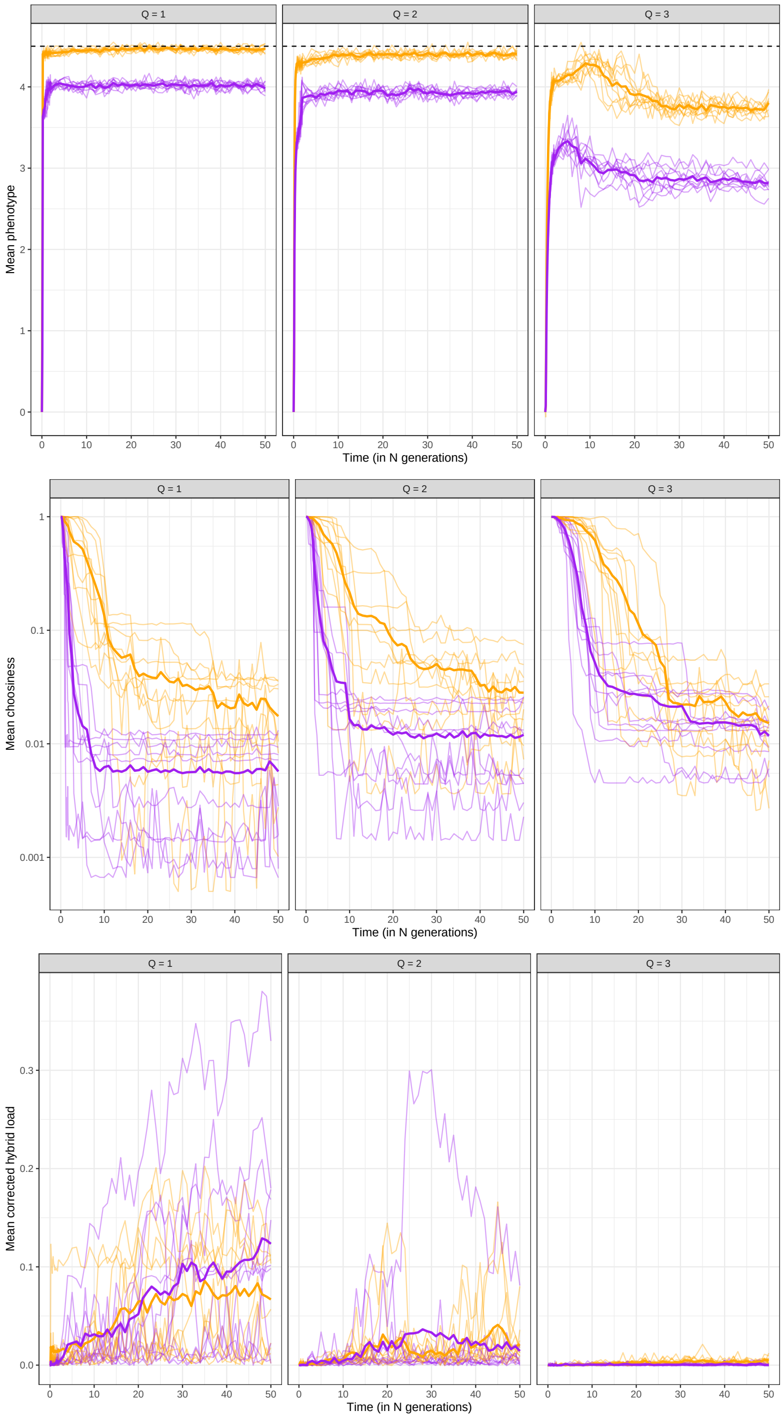


Figure S24: **Sharp fitness landscape (Q=1) promote the evolution of stronger RI barriers** Evolution of the phenotype (top), mate choice (middle; on a log scale) and intrinsic hybrid load (bottom) for different shape of the fitness landscape (given in the facet) and the two life cycles: "adult dispersal" in purple and "juvenile dispersal" in orange. Each thin line corresponds to a different replicate, and the thick line to the mean over the 10 replicates. For the top panel, the dashed line depicts the expected mean phenotypic values, given by $\bar{Z}_j = (1-m)Z_j^A + mZ_j^B$. The genetic map corresponds to the "default" map and other parameters correspond to the default value and are given in Table 1

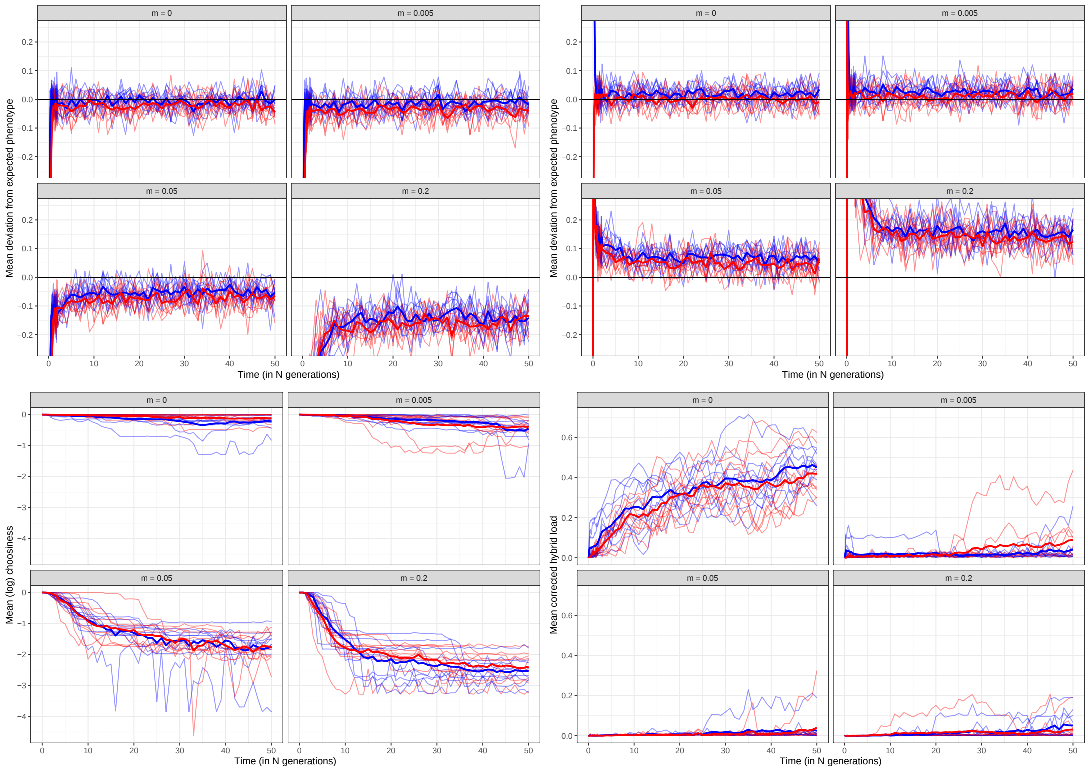


Figure S25: **Mutational bias has no effect when populations are locally adapted.** Evolution of the deviation of the phenotype from its expectation ($\bar{Z}_j = (1-m)Z_j^A + mZ_j^B$) (in environment A top left and in environment B top right), mate choice (bottom left; on a log scale) and intrinsic hybrid load (bottom right) depending on the position of the optimum in the fitness landscape. The ancestral phenotype is defined as $Z = \{0, 0, 0\}$. For the symmetric case displayed in blue, the optimal phenotypes in each environment are given by $Z = \{2.887, 2.887, 2.887\}$ and $Z = \{-2.887, -2.887, -2.887\}$ respectively. For the asymmetric case displayed in red, the optimal phenotypes in each environment are given by $Z = \{6.774, 6.774, 6.774\}$ and $Z = \{1, 1, 1\}$ respectively. In both case, the distance between optimum is therefore the same. Each thin line corresponds to a different replicate, and the thick line to the mean over the 10 replicates. For the phenotype, only the first trait is displayed. Other parameters correspond to the default value and are given in Table 1

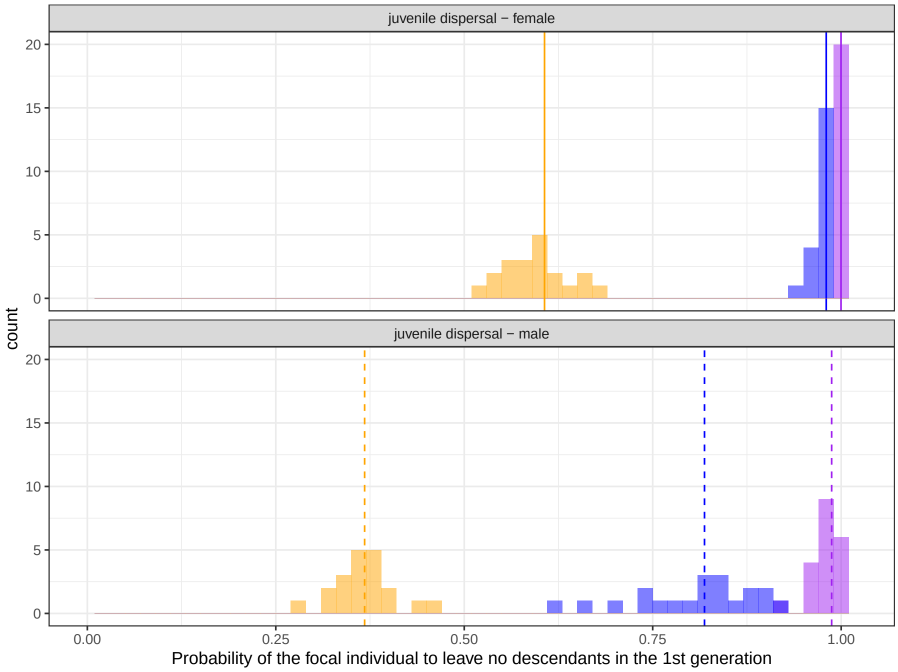


A/ Juvenile dispersal

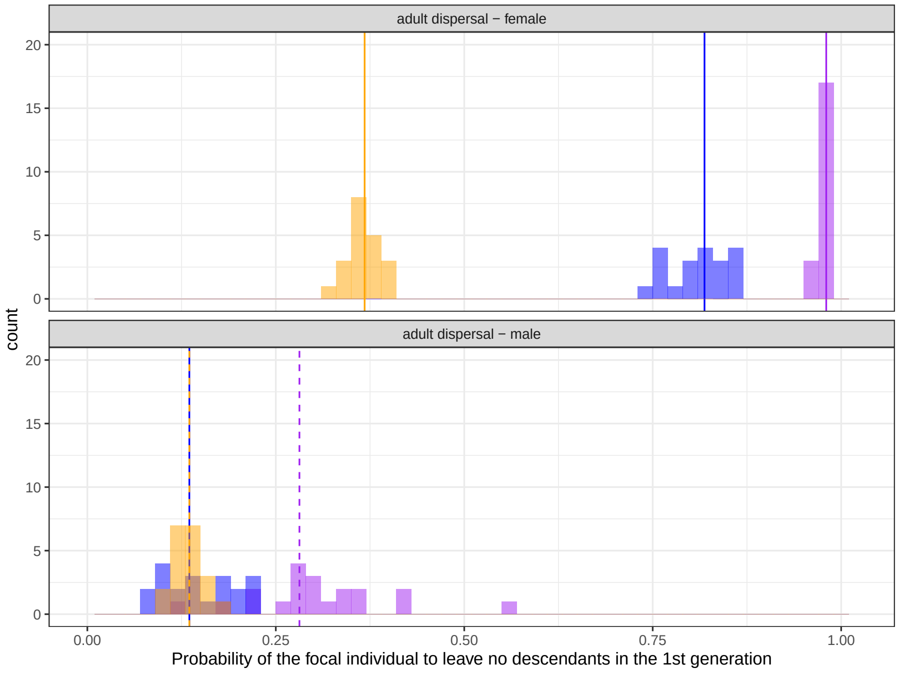


B/ Adult dispersal

Figure S26: **Analytical prediction for the probability of not leaving descendants in a single generations matches the simulations** Distribution of the proportion of focal individual that do not leave descendants in the first generation for the neutral marker, in the presence of the barriers to reproductive isolation. This distribution was computed for freely recombining marker for females and males and for both life cycles. Color indicates the strength of local adaptation and choosiness (orange $P_C = 0.5$, $w_f = 0.5$; blue $P_C = 0.1$, $w_f = 0.1$; purple $P_C = 0.01$, $w_f = 0.01$). The vertical line corresponds to the analytical prediction derived in the main text for males (solid line; eq. (A2) and females (dashed line; eq. (A3)). Here, we used population that were started with all barriers predefined and in isolation, to minimize the variance between replicates. Other parameters correspond to the the default values.

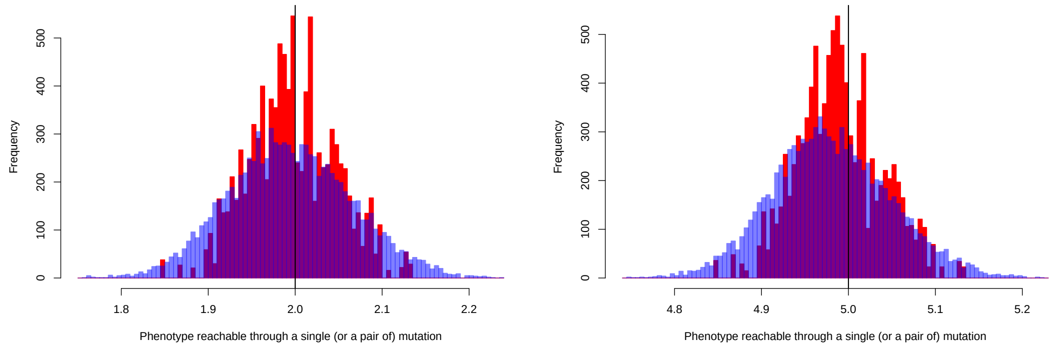


A/ Environmental phenotype, $Z_j^A = 2$

B/ Environmental phenotype, $Z_j^A = 5$

Figure S27: **Mutation bias tends to shift phenotypic values towards zero.** Distribution of phenotype reachable through a single (red) or two mutations (blue), assuming the phenotype is at the environmental optimum ($Z_j = Z_j^A$) and the underlying genotype is homozygous at all sites. The effect of the new mutations were evaluated as heterozygous. We used the "default map" and randomly generated 10,000 single mutations (or pairs of) on this map. For $Z_j^A = 2$, the mean mutational effect for a single mutation was $-0.003$ and the joint effect for a pair of mutation $-0.005$ for a pair. The proportion of single mutation increasing the value of phenotypic trait was 0.439 and 0.46 for a pair of mutations. For $Z_j^A = 5$, the mean mutational effect for a single mutation was $-0.009$ and the joint effect for a pair of mutation $-0.019$ for a pair. The proportion of single mutation increasing the value of phenotypic trait was 0.376 and 0.38 for a pair of mutations.
